# Mosaic vaccination: how distributing different vaccines across a population could improve epidemic control

**DOI:** 10.1101/2020.11.22.393124

**Authors:** David V. McLeod, Lindi M. Wahl, Nicole Mideo

## Abstract

Although vaccination has been remarkably effective against some pathogens, for others, rapid antigenic evolution results in vaccination conferring only weak and/or short-lived protection. Consequently, considerable effort has been invested in developing more evolutionarily robust vaccines, either by targeting highly conserved components of the pathogen (universal vaccines) or by including multiple immunological targets within a single vaccine (multi-epitope vaccines). An unexplored third possibility is to vaccinate individuals with one of a number of qualitatively different vaccines, creating a ‘mosaic’ of individual immunity in the population. Here we explore whether a mosaic vaccination strategy can deliver superior epidemiological outcomes to ‘conventional’ vaccination, in which all individuals receive the same vaccine. We suppose vaccine doses can be distributed between distinct vaccine ‘targets’ (e.g., different surface proteins against which an immune response can be generated) and/or immunologically-distinct variants at these targets (e.g., strains); the pathogen can undergo antigenic evolution at both targets. Using simple mathematical models, we show that mosaic vaccination often outperforms conventional vaccination, leading to fewer infected individuals, improved vaccine efficacy, and lower individual risks over the course of the epidemic.

## 1 Introduction

After hygiene, vaccines are arguably the greatest success story in public health to date. Vaccines are responsible for the eradication of smallpox in humans and rinderpest in livestock, and have driven substantial declines in the incidence of numerous childhood illnesses. Between 2010 and 2015, an estimated ten million lives were saved by vaccines [1]. While other control measures, like drugs, are failing in the face of pathogen evolution, vaccines seem comparatively robust [2, 3].

Yet vaccines are not immune to the challenges posed by pathogen evolution. As a result of either high mutation rates or existing standing variation, many pathogen populations harbour diversity in relevant immune-signalling sites. If that diversity translates to relatively weak immune responses against strains other than the one to which a host was previously exposed, then vaccines—often produced from one or a few target strains—will fail to offer broad protection. It is precisely this combination of limited cross-immunity (i.e, protection against different strains), resultant strong selection [4, 5], and rapid evolution that necessitates yearly updating of the composition of seasonal influenza vaccines, for example.

Efforts are ongoing to improve vaccination strategies against evolving threats like influenza [6]. In particular, the search for more highly conserved targets of immune responses may yet produce a universal vaccine that need not be updated in the face of antigenic evolution [7–10]. Yet the consequences of such vaccines, including for pathogen evolution, have not been fully elucidated [11]. An alternative strategy is a vaccine cocktail, designed to elicit immune responses against multiple targets (i.e., epitopes) with a single vaccine [11]; much like drug cocktails, these are expected to be more robust in the face of evolution [12]. Multi-strain vaccines offer a different kind of cocktail (e.g., eliciting immune responses against different variants of the same target), which has been broadly useful for limiting disease caused by some pathogens (e.g., pneumococcal vaccines [13]).

Here we investigate a different vaccination strategy by asking if and when vaccinating individuals with one of a number of qualitatively different vaccines—essentially a ‘cocktail’ at the population level— produces better epidemiological outcomes than a strategy that gives every individual the same vaccine. More accurately, in the vernacular of resistance evolution, this would be a ‘mosaic’strategy [12]. The epidemiological consequences of using a mosaic vaccination strategy, to our knowledge, have not been explored. We analyze a mathematical model to specifically ask, first, if vaccines with different targets existed, what would be the optimal way to use them? Second, we consider the scenario of having a set of vaccines that target the same immunologic site, but different genetic (and antigenic) variants, for which limited cross-immunity may exist. Finally, we ask which source of variation across vaccine doses—targets or variants—can produce the best outcomes when vaccine escape is either likely or rare. Overall, we find that conventional vaccine programs, in which all individuals receive identical vaccines, are often outperformed by mosaic strategies that deliberately seed variation in the host population.

## 2 The model

Consider a pathogen modelled in a standard SIR framework (see Sup. Info. 2). Infection leads to sufficiently broad and long lasting immunity such that once infected, hosts cannot be reinfected over the timescale under consideration. Let *R* denote the basic reproductive ratio of the pathogen in an unvaccinated population in which the density of susceptibles is normalized to 1.

The pathogen has two potential vaccine ‘targets’, *A* and *B*, each of which may exhibit antigenic variation. Biologically, these targets could be different surface proteins, or different epitopes on the same surface protein. We use ‘variants’ to refer to immunologically-distinct versions of these targets, and ‘strains’ to refer to pathogens harbouring different variants, such that *A_i_B_j_* denotes the pathogen strain with variant *i* at target *A* and variant *j* at target *B* (see Figure 1**a**). Antigenic space at each target is one dimensional [14], while antigenic change is cumulative and equally likely at either target. Thus if the initial pathogen strain is *A*_0_*B*_0_, then the next strain will be either *A*_1_*B*_0_ or *A*_0_*B*_1_ with equal probability. Antigenic change can either be generated by *de novo* mutation within the focal population, or it can be imported from an external source such as a reservoir animal population or a geographic region in which the pathogen is endemic (e.g., as for influenza A [15, 16]).

**Figure 1:**
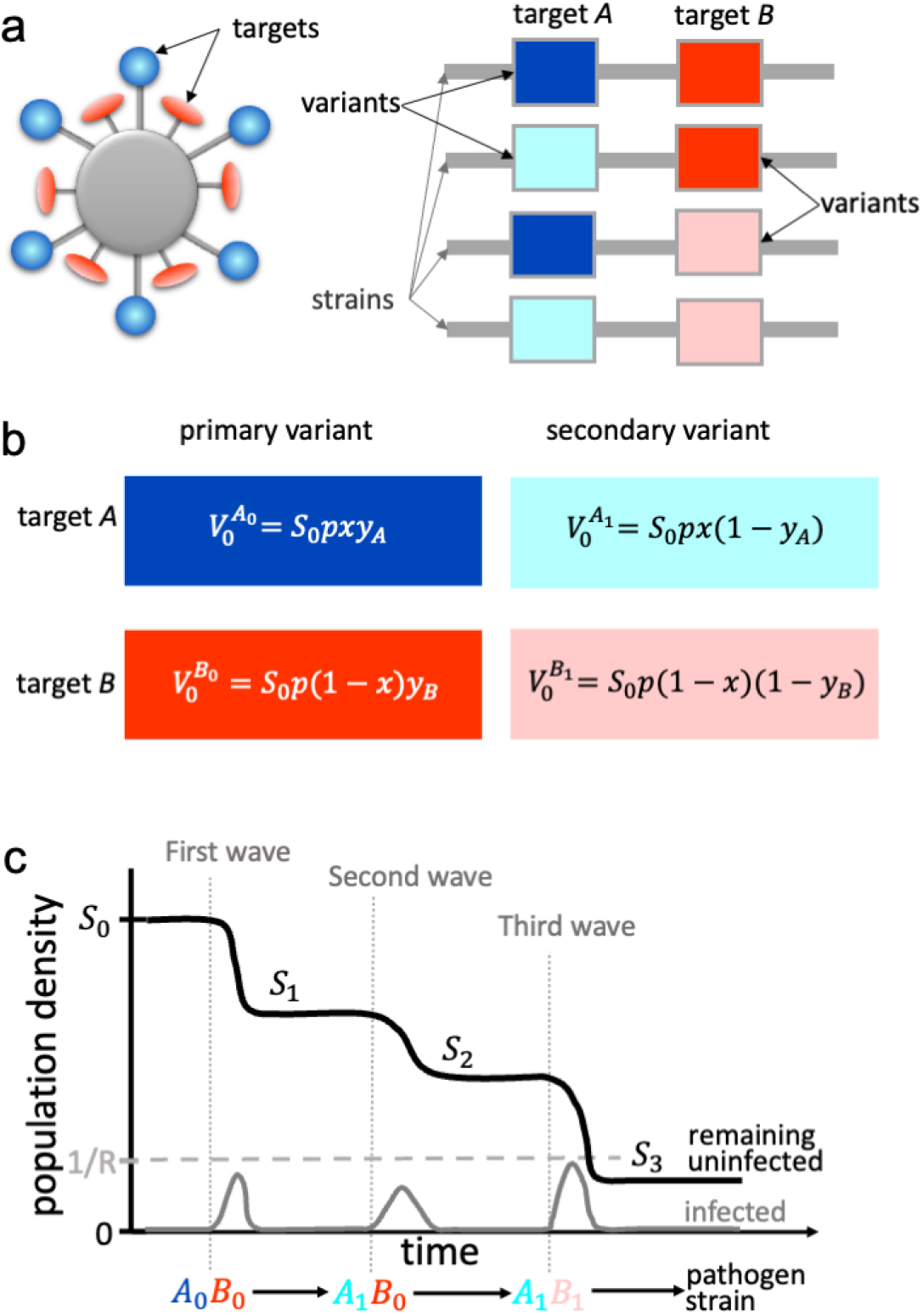
Model schematics. Panel **a** illustrates assumptions about pathogen structure (left) and genome (right); each line represents a unique pathogen genome, or ‘strain’, while the boxes represent specific loci. We use ‘targets’ to describe different surface proteins or, equally, different epitopes on the same surface protein. We use ‘variants’ to capture genetically (and more crucially, antigenically) distinct versions of a given antigenic target. Panel **b** gives the density of individuals vaccinated against different pathogen strains. A fraction *p* of all individuals in a population, *S*_0_, receive a vaccine. Vaccine doses are distributed across targets (a fraction, *x*, focus on target *A*, while 1 − *x* focus on target *B*) and variants (a fraction, *y_A_* or *y_B_*, focus on the primary variant of the given target). Panel **c** illustrates the typical dynamics of our model, where *S_n_* denotes the density of susceptible individuals (black curve) remaining after the *n*^th^ wave of the epidemic. The gray curve, indicating the density of infected individuals, captures our assumption that each wave reaches its epidemiological conclusion before another can begin, if the density of susceptibles is above 1/*R*. One possible epidemic sequence of strains is shown.

A fraction *p* of the population will be vaccinated, and the vaccine doses can be distributed amongst four possible candidates (Fig. 1**b**). A fraction *x* of the *p* doses are distributed to vaccine target *A*, and 1 − *x* to target *B*. Of the doses allocated to target *A*, a fraction *y_A_* are distributed to the most abundant variant in the population initially, *A*_0_, and 1 − *y_A_* to *A*_1_; likewise of the doses against *B*, a fraction *y_B_* will target *B*_0_, while 1 − *y_B_* target *B*_1_. While we restrict vaccine distribution to the set {*A*_0_, *A*_1_, *B*_0_, *B*_1_}, we make no such restriction for antigenic variation, and so it is possible that the variants *A_k_* and/or *B_k_, k* > 1 may emerge in the population.

Each vaccine reduces the probability of infection against its intended target/variant combination by a factor *χ*_0_, and more generally, reduces the probability of infection by a variant Δ*ℓ* antigenic units from its intended target/variant combination by a factor *χ*_Δ*ℓ*_. If vaccine cross-protection is sufficiently broad, the vaccine will not cause any antigenic evolution, and so the allocation of vaccine doses does not matter. As cross-protection decreases, however, the possibility that vaccination leads to antigenic evolution increases, and so how vaccine doses are distributed plays an increasingly prominent role. In our model, the broadness of vaccine cross-protection will depend upon both the units of antigenic space and the timescale of interest, and so we will focus upon the case in which vaccine cross-protection is negligible. This is done for simplicity, but we stress that the primary effect of broadening cross-protection is to reduce the impact of vaccine distribution.

We treat pathogen evolution as a sequence of discrete antigenic changes, each of which may (or may not) produce a “wave” of the epidemic, such that the overall infection process consists of a series of strain-specific waves (Fig. 1**c**). To simplify the mathematical analysis, we assume that these strain-specific waves occur sequentially, rather than simultaneously, i.e., a single strain dominates the infection process at any time. This assumption is valid provided antigenic change is sufficiently infrequent such that by the time a novel strain has risen to appreciable levels in the population, the epidemic wave by the previous strain has largely concluded.

Let *S*_0_ denote the initial density of individuals without infection-acquired immunity (susceptibles). *S*_1_ is then the density of susceptibles after the first wave (caused by the initial strain *A*_0_*B*_0_) and in general *S_n_*denotes the susceptible density following the *n*^th^wave. These individuals can be further divided based upon whether they have been vaccinated. Let *U_n_* denote the density of susceptible, unvaccinated individuals, and let 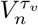 denote the density of susceptible individuals vaccinated against variant *υ* of target *τ*(*τ_υ_* ∈ {*A*_0_, *A*_1_, *B*_0_, *B*_1_}), following the *n*^th^ wave. Therefore,

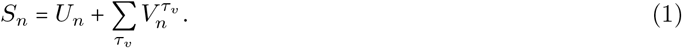

Suppose the initial densities are known, that is, *U*_0_ = *S*_0_(1 − *p*) and (for example) 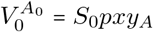. Then for a given sequence of antigenic changes, the state of the population after each wave of the epidemic could be determined by numerically integrating the full SIR model (Sup. Info. 2). By rescaling time, however, we can in fact directly compute the outcome of each wave using equations (2) and (3) below. First, we demonstrate in the Sup. Info. 3 that the remaining vaccinated individuals, after each wave, can be computed based on the sequence of remaining unvaccinated individuals. After the nth wave, the density of individuals vaccinated against *A*_1_ is:

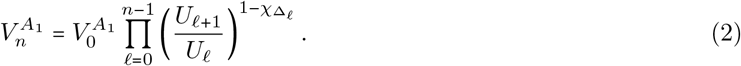

Here, Δ_*ℓ*_ measures any antigenic difference between *A*_1_ and the variant of A in the strain causing the *ℓ*^th^ wave, such that for example *χ*_Δ_*ℓ*__ = *χ*_0_ if the strain carries *A*_1_. The analogous variables, 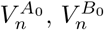, and 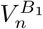 are computed similarly (Sup. Info. 3). In general, *U_n_*, 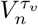, and *S_n_* will depend upon which of the 2^*n*−1^ sequences of antigenic changes occurred.

Given a particular sequence of *n* − 1 antigenic changes, from equations (1) and (2), the only unknowns are *U*_1_,…, *U_n_*. To compute the *U_i_*, we first determine whether the reproductive number of the pathogen strain with *n* antigenic changes is greater than one (Sup. Info. 3). If it is less than or equal to one, then *U_n_* = *U*_*n*−1_, otherwise *U_n_* is the solution of

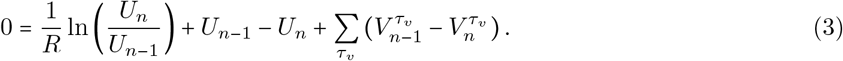

Once *U*_1_,…, *U_n_* are computed using equation (3), we can then calculate *S_n_* (using equation (1)).

### 2.1 The likelihood of vaccine escape

Given our assumption about the strength of immunity acquired through infection, there can be at most a single wave in an unvaccinated population. In a vaccinated population, however, antigenic change may cause successive waves as strains escape vaccine coverage. Our analysis will therefore focus upon two representative scenarios of vaccine escape; in both we are interested in the expected remaining uninfecteds, denoted 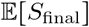. In the first scenario, we assume that vaccine escape is rare, i.e., at most one antigenic change might occur over the timescale of interest. Thus, following the initial potential wave by strain *A*_0_*B*_0_, with probability *ω* (for *ω* small) a single antigenic change occurs at either target with equal probability, while with probability 1 − *ω*, no antigenic change happens. Thus 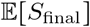 is the average of these possibilities (Sup. Info. 4).

In the second scenario, vaccine escape is common, and thus the sequence of epidemic waves is not limited by antigenic change. Specifically, following the initial potential wave by strain *A*_0_*B*_0_, the pathogen will undergo successive one unit antigenic changes (and successive potential waves) until the density of remaining individuals without infection-acquired immunity is less than or equal to 1/*R*; at this point, the population will have herd immunity and so any novel strains will be unable to cause an epidemic wave. In this case, 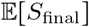 is the average remaining uninfected individuals computed over all possible antigenic sequences terminating in *S_n_* ≤ 1/*R* (Sup. Info. 4).

We will take maximizing 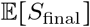 as the metric for evaluating the efficacy of particular vaccination strategies. An important point to note is that when vaccine escape is common, the best outcome we can hope to achieve is 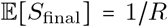. Thus irrespective of vaccine coverage, the population will ultimately be protected by herd immunity acquired through infection. The purpose of vaccination is then to ‘ease’ the population gradually down to the herd immunity threshold so that when it is reached, there are few active infections in the population and ‘overshoot’ (i.e., reducing the susceptible fraction to below 1/*R*) can be avoided [17]. (NB: Overshoot occurs when the herd immunity threshold is reached while many active infections exist, since even though, on average, infections are not able to replace themselves at this point, further transmissions occur as the wave declines.) In contrast, when vaccine escape is rare, considerably better outcomes than 1/*R* can be achieved.

Although we focus upon choosing vaccination strategies that maximize 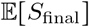, we are also interested in the consequences of these strategies for other metrics of vaccine success (Sup. Info. 6). The first metric of interest is vaccine efficacy, defined as

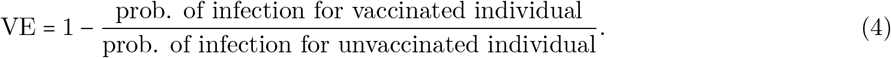

Here the probability of infection is computed over the entire epidemic. Vaccine efficacy measures the reduction in infection due to vaccination. Second, given an individual is both vaccinated and infected, what is the probability that they were infected by the strain they were vaccinated against (the ‘matched’ strain)? We refer to this as vaccine matching, calculated as

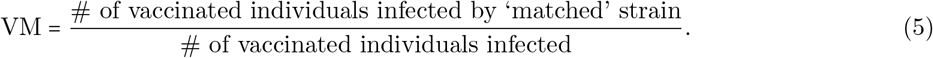

The importance of vaccine matching is that although our model assumes that the vaccine match determines only the probability of infection (i.e., χΔℓ), there may be additional impacts including on the severity of symptoms and, potentially, likelihood of hospitalization. Thus, all else being equal, a higher VM is desirable from a public health perspective.

### 2.2 Epidemic cases

To provide a baseline for comparison, consider ‘conventional’ vaccination, in which all doses are directed towards the primary variant of target *A*: *x* = 1 and *y_A_* = 1. Here it is helpful to divide the space of epidemic outcomes into three qualitatively different cases, based on the expected outcome of conventional vaccination with vaccination coverage, *p*, and vaccine strength, *χ*_0_ (Fig. 2).

**Figure 2:**
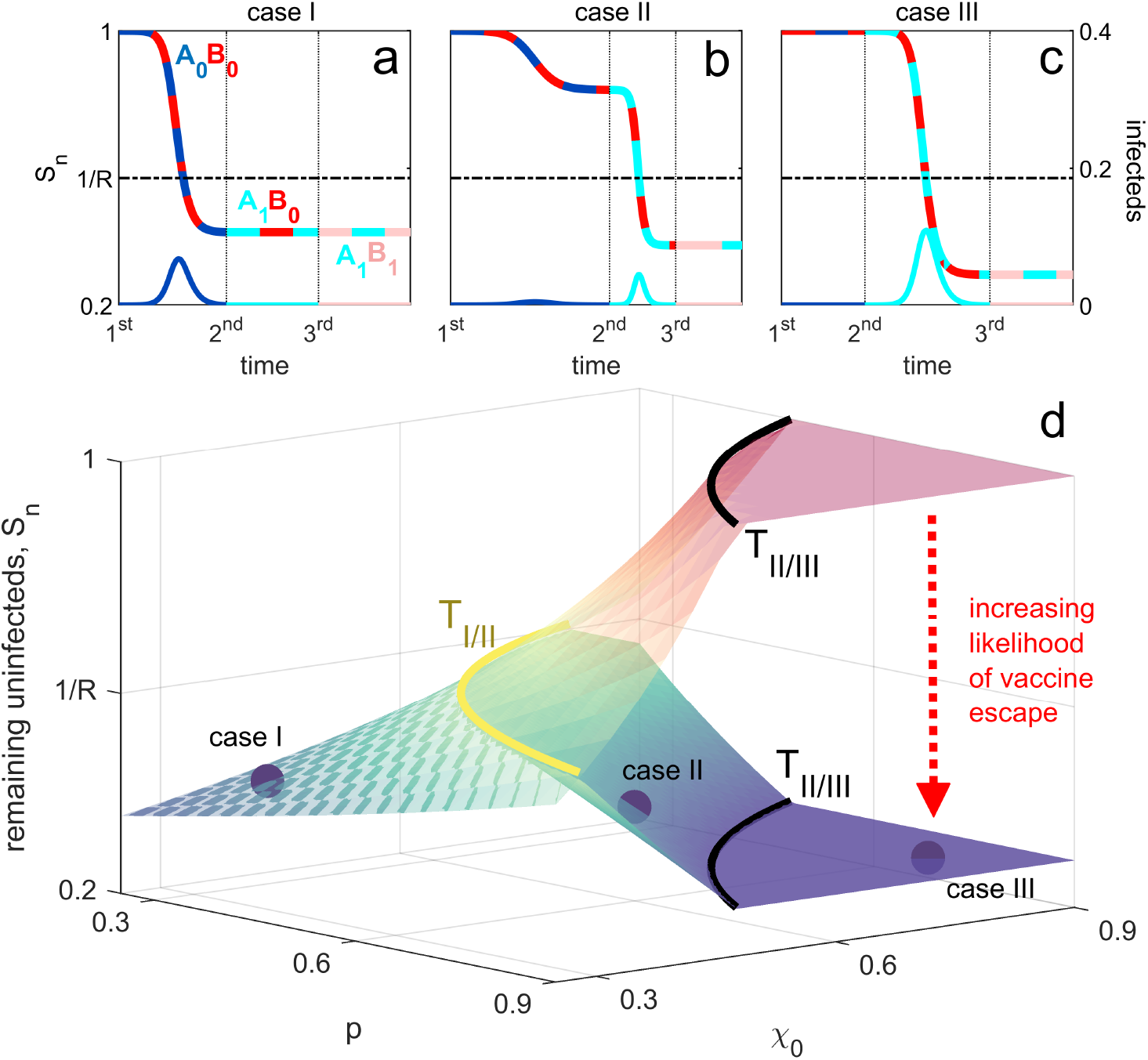
The three qualitatively different epidemic regimes under conventional vaccination. In case I, a single epidemic by the initial strain *A*_0_*B*_0_ occurs (panel **a**). In case II, there can be two epidemics, one by the initial strain, and one by the strain *A_i_B_j_*, here assumed to be *A*_1_*B*_0_ (panel **b**). In case III, the initial strain is blocked and a single epidemic wave occurs by a secondary (antigenically novel) strain (again, assumed here to be *A*_1_*B*_0_; panel **c**). In panel **d** we show the outcome in terms of the remaining uninfected individuals, *S_n_*, as we vary vaccine coverage, *p*, and vaccine strength, *χ*_0_, where the upper translucent surface is the outcome achieved when vaccine escape does not occur, whereas the lower surface is the outcome achieved when vaccine escape is common. The curves delineating the thresholds between cases, *T_I/II_*(*p*) and *T_II/III_*(*p*), are shown for reference, while panels **a-c** show examples of the epidemic dynamics for each case. On the remaining susceptible curves, the two colours indicate the strain circulating in the population, whereas on the infecteds curves, the single colour indicates what variant caused the epidemic wave.

As illustrated in Figure 2**a**, case I occurs when protection is sufficiently weak that the initial wave reduces the density of susceptibles to below 1/*R* and no further waves can occur; we denote the vaccine strength threshold at which this occurs as *T_I/II_*(*p*) (Sup. Info. 5). Thus when *χ*_0_ ≤ *T_I/II_*(*p*), infection-acquired immunity (due to the large number of individuals who were infected) protects the population against further waves (Fig. 2**a**).

Case II occurs when coverage is of intermediate strength. As shown in Figure 2**b**, in case II the wave caused by the initial strain is such that *S*_1_ > 1/*R*, and so antigenic change can generate subsequent waves until *S_n_* ≤ 1/*R* (Fig. 2**b**). Thus, in case II, antigenic change is possible but not necessary for an epidemic. We denote the vaccine strength threshold for the upper limit of case II as *T_II/III_*(*p*) (Sup. Info. 5), and thus case II occurs when *T_I/II_*(*p*) < *χ*_0_ ≤ *T_II/III_*(*p*).

Finally, in case III, antigenic change is necessary for an epidemic to occur. As illustrated in Figure 2**c**, in case III vaccine protection is sufficiently strong that the reproductive number of the initial strain is less than one and so antigenic change at target A must occur before the epidemic proceeds, *χ*_0_ >_*II/III*_(*p*). Thus the first wave can only be caused by a strain with variant *k* ≥ 1 at target A (Fig. 2**c**).

## 3 Results

### 3.1 Vaccines distributed across targets

First, we consider distributing equally-effective vaccines between two targets: a fraction *x* of available vaccine doses are used against target *A* and 1 − *x* are used against *B*. All doses are to be allocated to the primary variant at each target, that is, we will vary *x* while *y_A_* = *y_B_* = 1. Because both vaccines offer equal protection against the initial strain, the choice of *x* will only impact subsequent waves, to which conventional vaccination provides no protection (under our assumption of limited cross-immunity). As a result, mosaic vaccination (0 < *x* < 1) will be at least as good as conventional vaccination in terms of maximizing 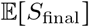. In fact, when antigenic evolution either can cause secondary waves (case II, Fig. 3**a**) or is necessary for an initial wave (case III, Fig. 3**b**), any variation in targets across vaccine doses, 0 < *x* < 1, will be superior to conventional vaccination in terms of maximizing the expected number of individuals who do not contract the disease.

**Figure 3:**
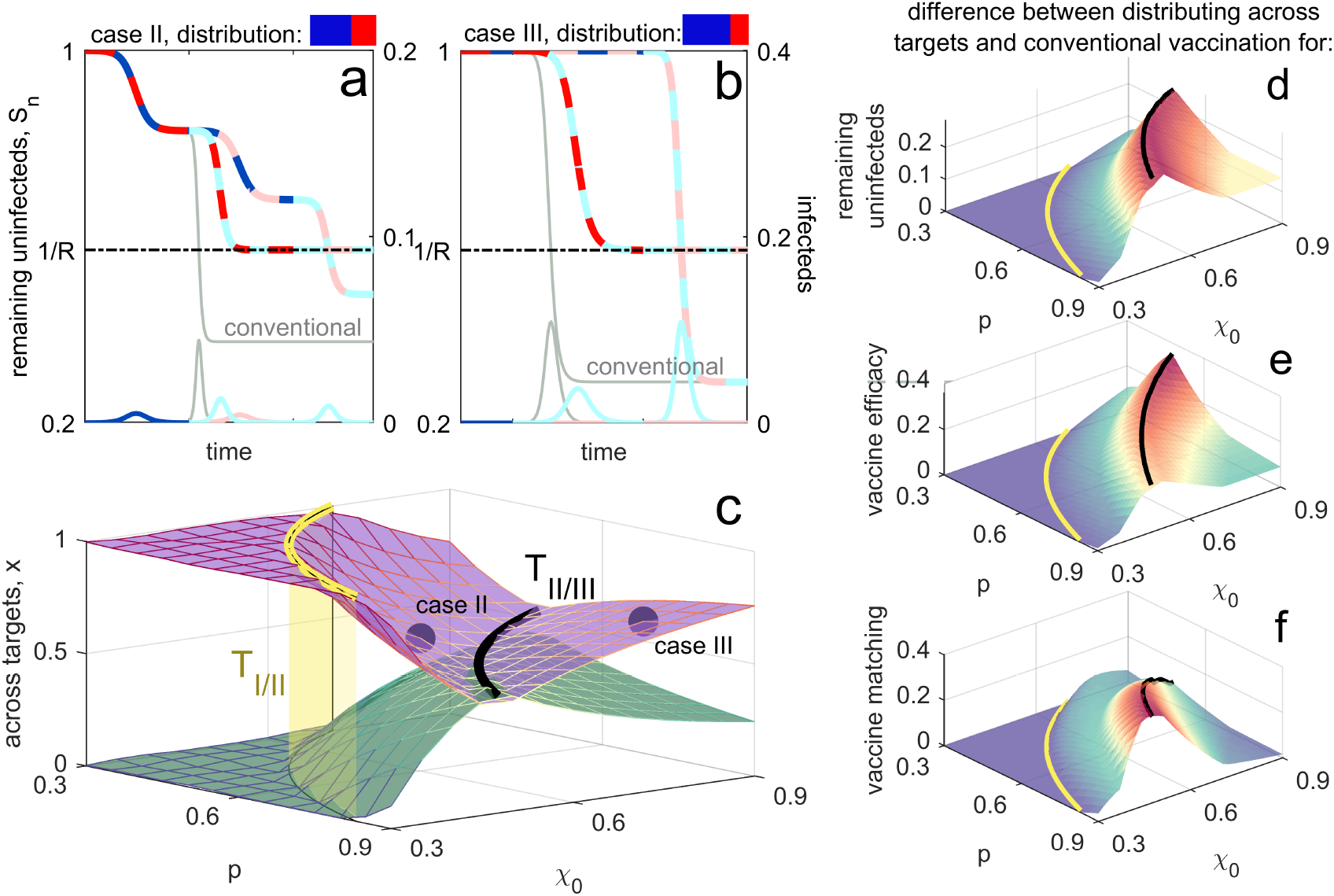
Optimal distribution of vaccines across targets when vaccine escape is common. Panels **a**,**b** show example epidemic dynamics for case II (panel **a**) and case III (panel **b**), with conventional vaccination (light grey) shown for reference; colours used are as in Figure 2. Values of *p, χ*_0_ and the optimal distribution *x*\* for these plots are indicated by the purple circles in panel **c**. Following the initial epidemic wave by strain *A*_0_*B*_0_, there are two possible sequences of interest, *A*_0_*B*_0_ → *A*_1_ *B*_0_ → *A*_1_*B*_1_ and *A*_0_*B*_0_ → *A*_0_*B*_1_ → *A*_1_*B*_1_; both are shown. On the remaining susceptible curves, the two colours indicate the strain circulating in the population, whereas on the infecteds curves, the single colour indicates what variant caused the epidemic wave. Panel **a** illustrates that mosaic vaccination is superior to conventional vaccination regardless of the sequence, whereas in **b**, mosaic vaccination is better on average and never worse. The optimal distribution across targets (shown in panel **c** as vaccine coverage, *p*, and strength, *χ*_0_, vary) maximises the remaining susceptibles averaged over these sequences; when *x* is optimal, so is 1 − *x* (purple, green). Panels **d-f** show how distributing across targets outperforms conventional vaccination for three metrics; positive values indicate the degree to which mosaic vaccination is superior. Panel **d** shows the difference in remaining uninfecteds, *S_n_*. Panel **e** shows the difference in vaccine efficacy, measured as attack rate over the course of the epidemic (Sup. Info. 7). Panel **f** shows the difference in vaccine matching, defined as the probability that an individual that is both vaccinated and infected is infected by a strain that they were vaccinated against. In all panels, the thresholds *T_I/II_*(*p*) (yellow surface) and *T_II/III_*(*p*) (black lines) are included for reference; beyond the yellow surface (i.e., further reducing *p* or *χ*_0_), vaccine efficacy is sufficiently low that evolution has no effect on the epidemic, and any distribution of vaccines between targets will produce the same outcome (for a given *p* and *χ*_0_).

Perhaps more surprisingly, Figure 3**c** demonstrates that for case II and III, an equal distribution of doses between targets, *x* = 1/2, is rarely the best strategy for maximizing 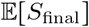, regardless of the likelihood of vaccine escape. This is because there are two equally probable sequences of antigenic change, differentiated by which target changes first (i.e., is the second wave caused by *A*_1_ *B*_0_ or *A*_0_ *B*_1_?). We can choose *x* to maximize vaccine protection against either one of these sequences. If vaccine escape is common, this means choosing *x* so that for one of the sequences, *S*_final_ = 1/*R* as shown in Fig. 3**a**,**b**. If vaccine escape is rare, this means choosing *x* to block the second wave of one sequence, while allocating the excess doses against the other target (Fig. 4**a**,**b**). In comparison with *x* = 1/2, either of these choices of *x*\* will reduce protection against the other sequence. However, for a wide range of parameter space, Fig. 3**d** demonstrates that the average outcome of *x*\* across the two possible sequences outperforms the expected outcome when *x* = 1/2, and always outperforms conventional vaccination (Fig. 3**c,d**; SI Fig. 1**c,d**).

**Figure 4:**
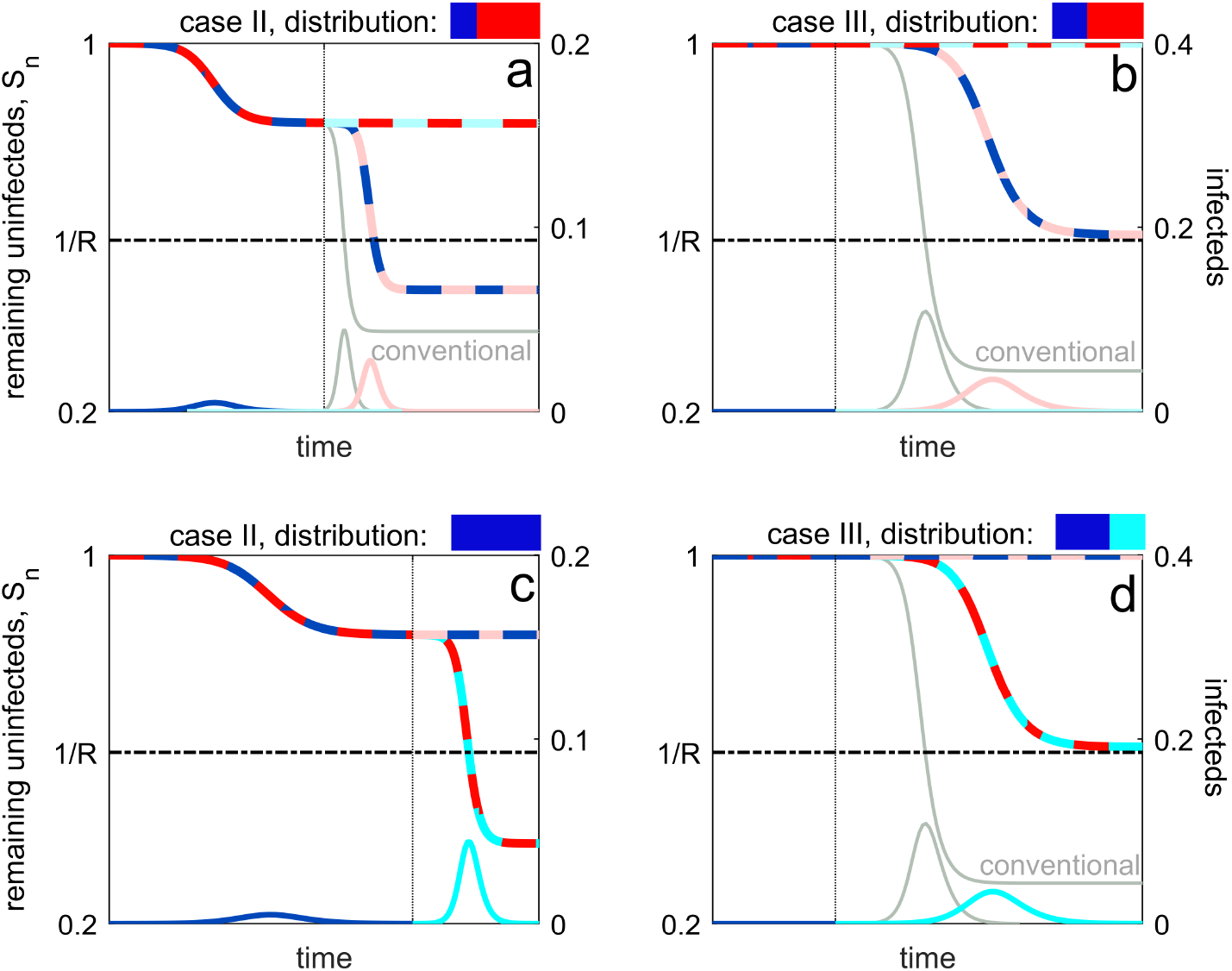
Epidemiological dynamics when optimally distributing across targets (panels **a, b**) or variants (panels **c, d**) when vaccine escape is rare. In all panels, there is a small probability of a single mutation at either target, giving rise to two equally probable sequences, *A*_0_*B*_0_ → *A*_1_*B*_0_ and *A*_0_*B*_0_ → *A*_0_*B*_1_; both of these sequences are shown. For the latter sequence, conventional vaccination blocks the second epidemic wave and so matches with the horizontal multi-coloured line in each panel. For the former sequence, conventional vaccination performs poorly, and is shown in light grey, except in panel **c** when the optimal distribution is also conventional vaccination. In Sup. Info. Figures 1 and 2, we show the optimal distribution and the performance metrics for mosaic vaccination when vaccine escape is rare.

The advantage of distributing across targets over conventional vaccination is not limited to increasing the remaining uninfecteds, 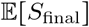.

Fig. 3**e** and **f** show that distributing across targets also increases vaccine efficacy and vaccine matching; results which also hold when vaccine escape is rare (SI Fig. 1**e**,**f**). Thus not only does mosaic vaccination improve population level outcomes, it also reduces individual-level risks associated with weaker protection against symptoms from mismatched vaccines.

### 3.2 Vaccines distributed across variants

Next, we consider distributing equally-effective vaccines between two variants of target *A*, that is, setting *x* = 1 and varying *y_A_*. Clearly, any *y_A_* < 1 will reduce the protection against the initial strain, and so if vaccine coverage is sufficiently weak (case I), all doses should be allocated to the primary variant, *y_A_* = 1 (conventional vaccination). In case II and III, however, conventional vaccination performs poorly because the population is effectively unvaccinated against any variation at target A (Fig. 2**b**,**c**).

If vaccine escape is rare, then it is optimal to maximize protection against the primary variant, *A*_0_. In case II, because conventional vaccination is not sufficiently strong to block the initial *A*_0_*B*_0_ epidemic wave, all doses should be allocated against *A*_0_ (*y_A_* = 1; Fig. 4**c**). In case III, because conventional vaccination is sufficiently strong to block the *A*_0_*B*_0_ epidemic wave, *y_A_* should be chosen such that the reproductive number for strain *A*_0_*B*_0_ is reduced to one. Here, any excess doses not needed to block the *A*_0_*B*_0_ epidemic wave are diverted to protect against the (rare) possibility that a strain carrying the variant *A*_1_ emerges (Fig. 4**d**; SI Fig. 2).

When vaccine escape is common, we can always choose *y_A_* to ensure that 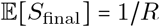, as illustrated in Fig. 5**a,b**). Thus in contrast to distributing vaccine doses between targets, when distributing across variants we can, in theory, always attain the optimum. This can be achieved with two values of *y_A_*: we can choose *y_A_* = *y*\*, which provides sufficiently weak protection against the initial strain such that *S*_1_ = 1/*R*; or we can choose *y_A_* = *y*^•^, where *y*^•^ > *y*\* such that only after antigenic change at target *A* will 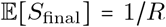 (Fig. 5**c**; Sup. Info. 8). Although both *y*\* and *y*^•^ maximize 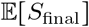, from a public health perspective *y*^•^ is the superior option as it maximizes 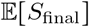 after at least one antigenic change, and so two potential waves, rather than one epidemic and no antigenic change. By staggering the waves, the peak burden on the healthcare system will be reduced (e.g., [18]). Indeed, the less desirable outcome of *y*\* could simply be achieved using conventional vaccination with artificially lowered coverage. In Figure 5**a,b** we show example epidemiological dynamics for *y*^•^.

As was the case when distributing across targets, the advantage of distributing across variants (using *y*^•^) over conventional vaccination is not limited to increasing the remaining uninfecteds, 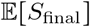 (Fig. 5**d**). Distributing across variants also increases vaccine efficacy (Fig. 5**e**) and vaccine matching (Fig. 5**f**). In case III, these results also hold when vaccine escape is rare (SI Fig. 2**e**,**f**). Thus distributing across variants improves both population level outcomes as well as reducing individual risk.

**Figure 5:**
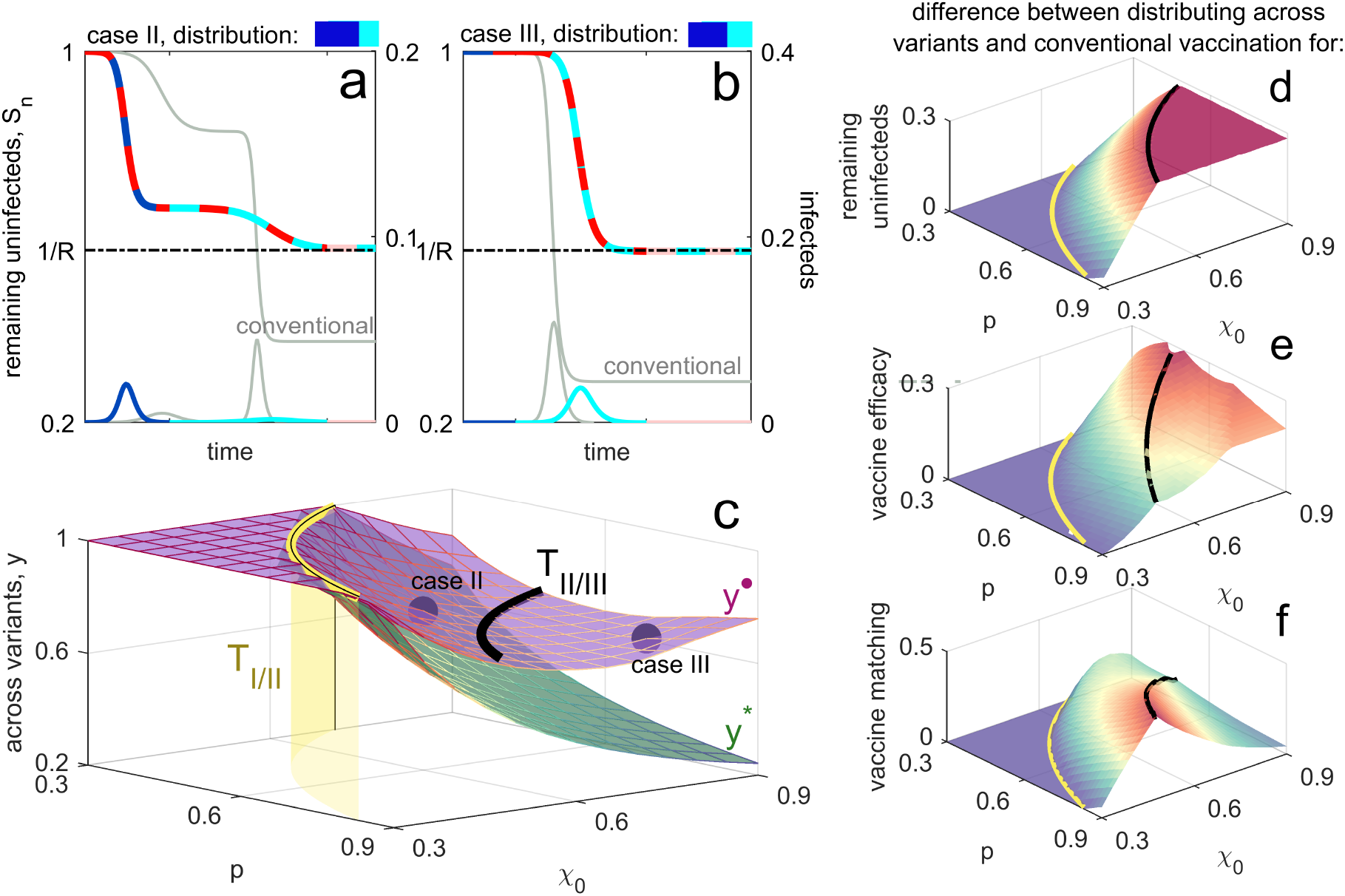
Optimal distribution of vaccines across variants when vaccine escape is common. Panels **a**,**b** show example epidemic dynamics for case II (panel **a**) and case III (panel **b**), with conventional vaccination (light grey) shown for reference. Values of *p, χ*_0_ and the optimal distribution *y*^•^ for these plots are indicated by the purple circles in panel **c**. For both panels, there is only one sequence of interest, corresponding to antigenic changes at target *A*: *A*_0_*B*_0_ → *A*_1_ *B*_0_ → *A*_2_*B*_0_. In case II, the optimal distribution weakens protection against the first epidemic wave, as compared to conventional vaccination so as to provide better protection against the second; the outcome across both epidemics is substantially improved. In case III, the optimal distribution across variants is to block the epidemic wave by strain *A*_0_*B*_0_ and then allocate any excess doses against variant *A*_1_. Panel **c** shows the optimal distribution across variants, *y*^•^ and *y*\* as vaccine coverage, *p*, and strength, *χ*_0_, vary. Panels **d-f** show how distributing across variants (using *y*^•^) outperforms conventional vaccination for three metrics; each panel shows the difference between distributing across targets and conventional vaccination. Panel **d**, shows the difference in remaining uninfecteds, 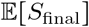. Panel **e** shows the difference in vaccine efficacy, measured as attack rate over the course of the epidemic. Panel **f** shows the difference in vaccine matching, defined as the probability that an individual that is both vaccinated and infected is infected by a strain that they were vaccinated against. In all panels, the thresholds *T_I/II_*(*p*) (yellow surface) and *T_II/III_*(*p*) (black lines) are included for reference.

### 3.3 Distributing across targets and variants simultaneously

Finally, consider distributing vaccines across both targets and variants. When we are free to choose all three of (*x, y_A_, y_B_*), many different combinations can lead to similar *S_n_*, so we can apply further constraints to identify optimal combinations (Sup. Info. 9). We opt to impose the constraint that *y_A_* = *y_B_* = *y*; this allows us to isolate whether distributing over targets or variants is more important.

Under these circumstances, the outcome is as we would expect from consideration of our previous results. When the likelihood of vaccine escape is low, the optimal solution is to rely upon distributing doses between targets (varying *x*) while targeting the primary variant, *y* = 1. As vaccine escape becomes more common, the optimal strategy is to distribute between variants (varying *y*), while keeping doses equally divided across targets, *x* = 1/2 (Sup. Info. 9). The reason for this is intuitive: when there is a low probability of a second wave, we want to maintain maximum protection against the initial strain (*y* = 1), but we are free to distribute between targets so as to block one of the epidemic sequences. As the likelihood of vaccine escape increases, a wave by a secondary variant is increasingly likely, and so we are more willing to sacrifice protection against the initial strain to allocate doses against the secondary variants. Although we previously showed *x* = 1/2 is not optimal when distributing across targets only, if we are free to distribute across both variants and targets, and are allocating doses to the both the primary and secondary variant, the logic changes. Specifically, since either epidemic sequence is equally likely, when relying upon distributing between variants it makes sense to distribute equally between targets (*x* = 1/2; SI Fig. 3).

## 4 Discussion

We have used a simple model to explore the optimal distribution of vaccines that differ in antigenic targets and/or specific variants at those targets (i.e., strains). We show that with weak vaccines (either low coverage or strength), evolutionary change in the pathogen population does not alter epidemiological outcomes and so a mosaic vaccination strategy, in which individuals receive one of a set of possible vaccines, is not better than a conventional strategy. In cases where pathogen evolution can lead to successive waves of an epidemic, or is required for an initial wave, using a mosaic vaccination strategy often leads to better outcomes than giving all individuals identical vaccines.

Vaccines distributed across targets are equally effective against the initial circulating strain, thus producing outcomes at least as good as conventional vaccination. However, if antigenic evolution occurs, the benefits of mosaic vaccination are realized. While a conventional vaccine would offer no protection against a strain that had evolved at one of the two targets (given our assumptions about cross-immunity), some protection would be maintained in a population that received a mosaic strategy. Perhaps surprisingly, the optimal distribution of vaccine doses between targets is not 50:50—better outcomes can be achieved on average with unequal allocation by providing greater protection against one sequence of antigenic changes, while still retaining some protection against the other possible sequence (unlike conventional vaccination).

While not our focus or motivation, it is interesting to consider our results in the context of the ongoing COVID-19 pandemic, given the rapid and parallel development of dozens of vaccines against SARS-CoV-2 [19–21]. While many COVID-19 vaccines in development target the same spike protein, some target epitopes in other genes [20] and the potential for multi-epitope (i.e., cocktail) vaccines is being investigated (e.g., [22–25]). If in the future multiple COVID-19 vaccines were approved with similar efficacies (i.e., roughly equal protection against the predominant circulating strain), then our results suggest that to minimize the risk of evolutionary escape and reduce the total number of individuals who get infected, the vaccines should be used in a mosaic strategy (*sensu* [12]). Multiple plausible vaccine targets exist for other infectious diseases (e.g., hemagglutinin (HA) and neuraminidase (NA) in influenza [26]), but whether each target would induce equal efficacy protection and is subject to an equal likelihood of mutation remain open questions (though see [5] for similar estimates of non-synonymous mutation rates in HA and NA within hosts). Our model provides a framework for building in these details and evaluating the efficacy of mosaic vaccination strategies for specific diseases.

What happens when vaccines are distributed across variants of a given target is perhaps more subtle. If evolutionary change in antigens is unlikely, then diverting any vaccine doses away from the predominant strain can lead to worse epidemiological outcomes. However, even if antigenic change is unlikely, if vaccine efficacy (*χ*_0_) is sufficiently strong or coverage (*p*) is sufficiently high (i.e., our case III), then from a population perspective better outcomes can be achieved by vaccinating a fraction of the population against a secondary variant. As the likelihood of waves from antigenically-distinct strains increases, then a trade-off emerges between protection in an initial wave versus subsequent ones. Put simply, a small initial wave due to strong vaccine protection against an initial variant leaves a large pool of susceptible individuals that could be exploited by a subsequent strain (assuming weak cross-immunity from vaccination), while a large initial wave due to poor vaccine protection leaves few individuals susceptible to future waves (assuming strong cross-immunity from natural infections). The optimal strategy essentially titrates between these scenarios, offering intermediate protection against initial and subsequent strains and dampening—but not eliminating— individual waves. This intermediate strength optimal strategy echoes results from previous work: in an explicit two-strain epidemiological model, if vaccination disproportionately impacts one strain (and vaccine-induced immunity is weaker than natural immunity), then increasing vaccine coverage or strength can lead to outgrowth of infections with the second strain and worse outcomes overall [27, 28].

Distributing across variants seems intuitively risky. For influenza, for example, substantial work goes into choosing which variant of each flu subtype will be included in a vaccine in a given year (e.g., [29]). Part of the decision is based on the frequency of circulating variants and the likelihood that any one variant will seed the coming year’s seasonal epidemic. If there is good reason to expect a particular variant will circulate predominantly, it would seem unethical to allocate any vaccine doses to a different variant. Yet our results show that over a considerable range of parameters (i.e., vaccine efficacy and coverage, likelihood of antigenic change), the conventional strategy does not give rise to the best outcomes and so the principle of equipoise would not be breached by distributing vaccines across variants. Of course, our model assumes that the second, dominant strain to circulate in a given epidemic can be predicted. For influenza, evolutionary predictions are improving (e.g., [29–31]); our results suggest a novel way of using those predictions for vaccine design, assuming any practical barriers can be overcome.

Somewhat akin to mosaic vaccination, previous work has explored the epidemiological and evolutionary consequences of vaccines that produce variable effects across hosts. First, using a data-driven model of a viral disease of fish, Langwig et al. [32] showed that vaccines which induce variation in susceptibility across hosts can lead to better epidemiological outcomes than vaccines that have individually-invariant effects (analogous to theoretical results exploring the effects of natural variation in susceptibility [33–35]). This is because with variation, average susceptibility declines over the course of the epidemic, since hosts that are the most susceptible get infected earlier on average. Likewise, in our model, distributing vaccine doses across targets or variants changes the landscape of host susceptibility both initially and as circulating pathogens evolve. In our case, distributing across targets mitigates the increase in susceptibility that would occur with conventional vaccines following antigenic evolution, while distributing across variants can actually reverse it. Second, as an explanation for why few vaccines have failed in the face of pathogen evolution, relative to the alarming rise and spread of drug resistance, Kennedy & Read [2] argue that individual variation in response to vaccination, e.g., generating immunity against different antigens, may lead to more diverse selection pressures acting on pathogens. In practice, for influenza at least, there is strong evidence that individuals do respond qualitatively differently to the same vaccine, due to differences in past history of exposure (e.g., [36]; reviewed in [37]). Our work reinforces the idea that, in some cases, this variation can be beneficial from a public health perspective [2] and so generating such variation could be an explicit aim of vaccination.

Our model makes a number of key assumptions (Sup. Info. 10). First, antigenic change was assumed to arise without consideration of its source. This is reasonable if antigenic novelty originates outside the focal population. For example, it may come from a reservoir animal population or be otherwise imported from a source population (e.g., influenza A [15, 16]). If instead we focused strictly on antigenic change from de novo mutation in the focal population, then unless mutations are very likely, conventional vaccination tends to perform better than our results show; in this case it is typically better to ‘hit hard’ in the hopes of preventing antigenic change. Second, we assumed cross-protection was limited. The ‘broadness’ of vaccine cross-immunity determines the reduction in vaccine protection against new strains. If vaccine cross-immunity is broad, antigenic change will cause a small reduction in vaccine protection, and so a smaller subsequent wave. Thus increasing cross-protection increases the efficacy of conventional vaccination in case II and III. In our model, the broadness of cross-immunity will ultimately depend upon the units of antigenic space and the time scale of interest. Third, we assumed each wave consists of a single strain. Although this may be reasonable if antigenic variation originates elsewhere, it is less likely when variation is generated by de novo mutation. If our results were extended to include multi-strain waves, the predicted efficacy of conventional vaccination would be weakened, since whenever an escape mutant arises during an ongoing wave, there are more individuals without infection-acquired immunity and so available to be infected by the escape mutant. This would lead to a larger wave by the strain with limited vaccine protection.

While the evolutionary and epidemiological consequences of universal vaccines have received some theoretical attention (e.g., [38]), here we explored the potential for vaccination strategies to essentially generate universal coverage at the host population level by delivering variable vaccines to individuals. This mosaic strategy generates heterogeneity in the host population (and, thus, the fitness landscape for pathogens), which has long been thought to be protective against disease outbreaks [39]. Theory predicts that ‘naturally’ variable host populations are less likely to experience sustained disease spread (e.g., [34, 35]), and the protective effect of host variation has been empirically demonstrated in a number of experimental (e.g., [40, 41]) as well as natural systems [42], especially those in which rapid host evolution is unlikely [43]. Our work suggests that vaccination strategies that harness—and, in fact, generate—variation can often outperform conventional vaccines.

## Acknowledgements

This work was supported by a Research in Teams (RiT) grant through the School of Mathematical and Statistical Sciences, Western University, and by the Natural Sciences and Engineering Research Council of Canada (grants RGPIN-2019-06294 to LMW, RGPIN-2018-06017 to NM, and a postdoctoral fellowship to DVM). We thank the participants of the RiT workshop for discussion, Evan Mitchell and Josh LeClair for valuable suggestions, and Mallika Makkar for inspiration.

## Supplementary Information

### 1 Preliminaries

We consider a pathogen with two vaccine targets, *A* and *B*, each of which can exhibit antigenic variation. We let *A_i_B_j_* denote a pathogen strain with variant *i* at target *A* and variant *υ* at target *B*. At each target, antigenic space is one dimensional, such that as *i* (or *j*) increases, the strain *A_i_B_j_* is further away from the reference strain, which we denote *A*_0_*B*_0_. Using this reference strain, we will refer to the variant indexed 0 as the ‘primary’ variant and the variant indexed 1 as the ‘secondary’ variant, e.g., strain *A*_1_*B*_0_ has the secondary variant for target *A* and primary variant for target *B*.

Each vaccine has efficacy *χ*_0_ against its intended target/variant combination; for example, the probability that strain *A*_0_*B*_1_ infects individuals vaccinated against *B*_1_ is reduced by a factor 1 − *χ*_0_. More generally, a vaccine protecting against variant *j* = 0,1,… of one target reduces the probability of infection by a strain with variant *i* of the same target by a factor *χ*_|*i-j*|_. Thus unlike in the main text, here we allow vaccines to confer cross-protection, where the degree of cross-protection depends only on the antigenic distance between variants at the vaccine target and not the variant at the other target. In contrast, infection-acquired immunity offers full protection against all strains, thus vaccine protection is inferior to infection-acquired immunity. We summarize our main results in Table 1.

**Table 1:** Summary of results. *S* denotes the remaining uninfected individuals; 1/*R* denotes the herd immunity threshold.

| | **between targets**<br>$A_0 : B_0$ | **between variants**<br>$A_0 : A_1$ |
| --- | --- | --- |
| **Case I:** even with vaccination, $A_0 B_0$ wave is severe and reduces $S$ to below $1/R$. | all allocations equally effective | use $A_0$ only |
| **Case II:** vaccination is strong enough that $A_0 B_0$ wave leaves $S$ above $1/R$.<br>**Case III:** vaccination is strong enough that $A_0 B_0$ wave is prevented. | **Cases II and III:** mosaic is always at least as good as (and often better than) conventional (details below) | **Cases II and III:** mosaic can be better than conventional; outcome depends on escape probability (details below) |
| Cases II and III, escape rare (escape with probability $\omega$) | A:B ratio has no effect on first wave. Allocate to block one of the two possible second waves. Average outcome outperforms 50:50 allocation, and is always better than conventional. | If escape is very rare, maximize protection against $A_0$: in Case II, allocate all doses to $A_0$; in Case III, allocate sufficient doses to block first wave, with extra doses toward $A_1$. As escape becomes more likely, balance shifts to protecting against $A_1$. See box below. |
| Cases II and III, escape common (best outcome is $S = 1/R$) | Similar to above, allocate so that one of the possible wave sequences ends with $S = 1/R$. Although this makes things worse than the 50:50 outcome if the other mutation happens first, the average outcome outperforms 50:50 allocation, which outperforms conventional. | Unlike conventional, mosaic vaccination can always achieve $S = 1/R$ in this case, after only one wave, or after two. To flatten the peak number of infections, allocate so that $S = 1/R$ after the second wave. |

### 2 Single epidemic wave

To model the epidemiological dynamics of a single epidemic wave, we use a standard SIR model with a single strain, say *A_ℓ_B_m_*, present. Let *β* be a rate constant reflecting the number of disease transmissions caused by a single infectious individual, per unit time, in a fully susceptible population. Infections have mean duration *δ* such that in the absence of vaccination, *R* = *βδ* reflects the basic reproductive ratio of this pathogen in a population where the density of susceptibles is normalized to 1.

Let *I_i_*(*t*) denote the density of infected individuals with strain *i* = *A_ℓ_B_m_* at time *t*. Let *U*(*t*) denote the density of unvaccinated individuals in the population at time *t*, and *V^τ_υ_^*(*t*) denote the density of individuals vaccinated against target *τ* and variant *υ* (*τ_υ_* ∈ {*A*_0_, *A*_1_, *B*_0_, *B*_1_}). Then the epidemiological dynamics are given by the system of ordinary differential equations

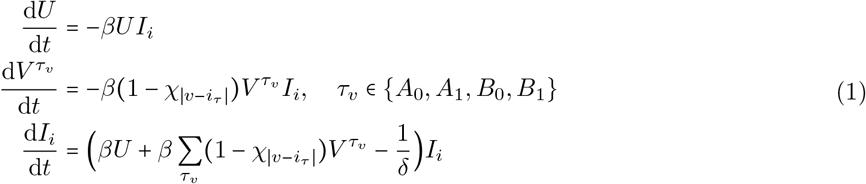

where *i_τ_* is the variant of strain *i* = *A_ℓ_B_m_* at target *τ*, i.e., *i_A_* = *ℓ* and *i_B_* = *m*, and so |*υ* – *i_τ_*| is the distance in antigenic space between the variant targeted by the vaccine, *υ*, and the variant carried by strain *i*. From system (1), the basic reproductive number of pathogen strain *i* is

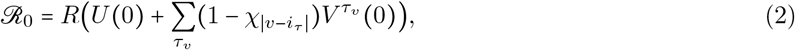

where *R* ≡ *βδ*. It follows that if 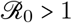, then an initially rare strain *i* will cause an epidemic.

The density of individuals who have not been infected by time *t* is

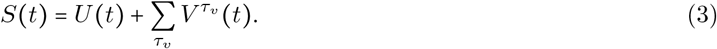

Our objective is to derive from system (1) an expression for the remaining uninfected individuals, *S*(*t*), after sufficient time has elapsed for the epidemic wave to conclude, that is, we are interested in *S*(∞). To do so, notice in system (1) that *U* is a monotonically decreasing function of time. Therefore, we can rescale time by *U*, which will give the reduced system

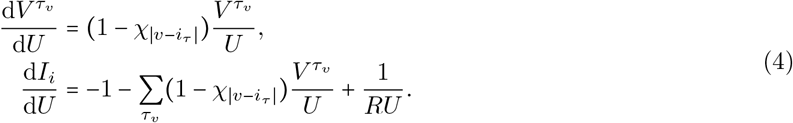

From (4), we can directly solve for *V^τ_υ_^*, that is,

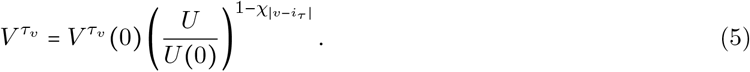

Using this result, we are left with the differential equation

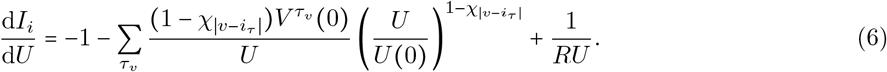

Applying the condition that strain *i* is initially rare, that is, *I_i_*(0) ≈ 0, this has solution

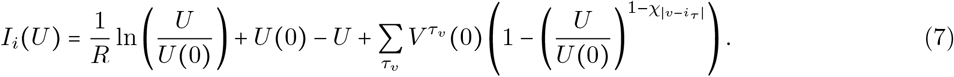

As *t* → ∞, we know that *I_i_* → 0, that is, the wave of strain *i* will eventually burn itself out of the host population. Therefore *U*(∞) is the solution of

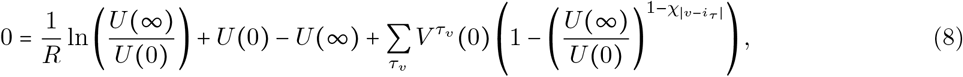

which can be written, using equation (5), as

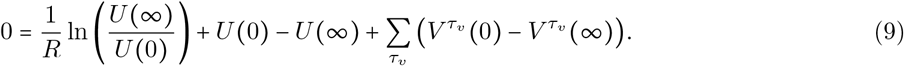

Therefore, there are two possibilities: if 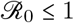, *U*(∞) = *U*(0), otherwise *U*(∞) is the solution of equation (9) satisfying 0 < *U*(∞) < *U*(0). Once we have *U*(∞), we can then substitute it into equation (3) to obtain *S*(∞).

### 3 Successive epidemic waves

We now extend equation (9) to consider a sequence of successive epidemic waves starting from the initial strain *A*_0_*B*_0_. To do so, let *U_n_* and 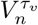 denote the density of susceptible, unvaccinated individuals and susceptible individuals vaccinated against *τ_υ_*, following the *n*^th^ epidemic wave. This epidemic wave will have been due to a strain *n* – 1 antigenic changes from the initial strain. Before any epidemic waves (or antigenic changes), i.e., *n* = 0, we have *S*_0_ susceptible individuals without infection-acquired immunity. We vaccinate a fraction *p* of these. Of the vaccinated individuals, a fraction *x* vaccine doses are directed towards target *A* and 1 − *x* directed towards target *B*. A fraction *y_A_* of the doses for target *A* are used against the primary variant, *A*_0_, while the remaining 1 − *y_A_* are used against the secondary variant, *A*_1_, likewise, a fraction *y_B_* of target *B* doses are used against *B*_0_, while the remaining 1 − *y_B_* are used against the secondary variant, *B*_1_. Therefore we initially have a density of *U*_0_ = *S*_0_ (1 − *p*) unvaccinated, susceptible individuals, while the initial density of individuals vaccinated against each target and variant combination is:

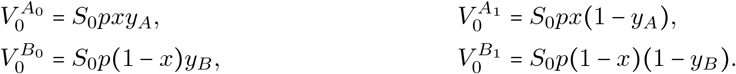

After the *n*^th^ epidemic wave, the next pathogen strain, *A_ℓ_B_m_*, will have undergone *n* = ℓ + *m* antigenic changes. There are 2^*n*^ possible antigenic sequences and *n* + 1 possible *A_ℓ_B_m_* strains. For a given sequence of antigenic changes culminating in strain *A_ℓ_B_m_* following the *n*^th^ epidemic wave, the basic reproductive number of strain *A_ℓ_B_m_* is

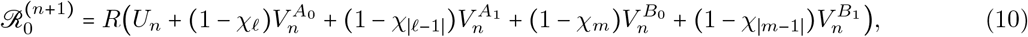

that is, 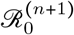 indicates whether or not the (*n* + 1)^th^ epidemic wave will occur. Using the sequence *A*_0_*B*_0_ → *A*_0_*B*_1_ → *A*_1_*B*_1_ as an example, the basic reproductive number of strain *A*_1_*B*_1_ is:

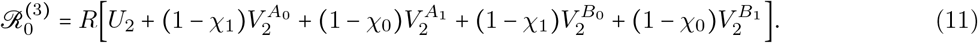

Note that, in general, *U_n_* and 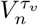 will be specific to the sequence of antigenic changes.

If 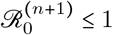, then *U*_*n*+1_ = *U_n_*, otherwise from equation (9), *U*_*n*+1_ is the solution of

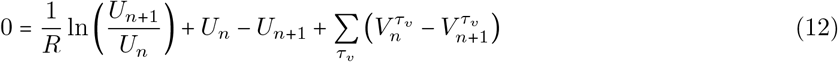

satisfying 0 < *U*_*n*+1_ < *U_n_*. Then using Equation (5) we have

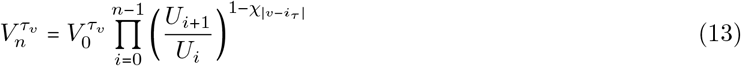

where *i_τ_* is the variant at target *τ* of the strain causing epidemic wave *i*, and

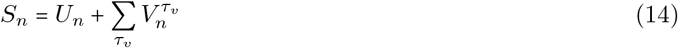

is the remaining uninfected individuals after the *n*^th^ epidemic wave. We are therefore equipped to compute the remaining uninfected individuals after the *n*^th^ epidemic wave for a given sequence of antigenic changes.

Although our derivations have made no assumptions about the ‘broadness’ of cross-immunity, in what follows we will focus upon the situation in which cross-immunity is limited. We discuss the implications of the ‘broadness’ of cross-immunity in more depth in Section 10.

### 4 The likelihood of vaccine escape

Because the outcome, *S_n_*, will depend upon the sequence of antigenic changes, in the main text we focus upon comparing the expected remaining uninfecteds, 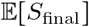 for two different scenarios. Specifically, the scenarios we consider are (i) when vaccine escape is rare, and (ii) when vaccine escape is common.

When vaccine escape is rare, we assume that following each epidemic wave, with probability *ω* a single antigenic change occurs at either target with equal probability, while with probability 1 − *ω*, no antigenic change happens. We assume that *ω* is sufficiently small, that is, vaccine escape is sufficiently rare, such that we can neglect the probability of more than one antigenic change over the time scale of interest. Therefore if we let *S*_2_(*A_j_B_k_*) denote the density of hosts without infection-acquired immunity following a second wave by strain *A_j_B_k_*, then 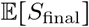 is given by

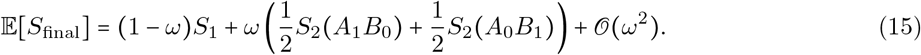

When vaccine escape is common, following the initial potential wave by strain *A*_0_*B*_0_, the pathogen will undergo successive one unit antigenic changes (and successive potential waves) until the density of remaining individuals without infection-acquired immunity is less than or equal to 1/*R*; at this point, the population will have herd immunity and so any novel strains will be unable to cause an epidemic wave. In this case, 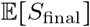 is the average remaining uninfected individuals computed over all possible antigenic sequences terminating in *S_n_* ≤ 1/*R*. In general this threshold will be reached within 1 or 2 antigenic changes (under the assumption of limited cross-protection, *χ*_|*z*|_ ≈ 0 for *z* > 0). For epidemics in which vaccine escape is common, we restrict our attention to situations in which we either distribute vaccines between targets, but not variants (so vary *x* while fixing *y_τ_* = 1), or between variants, but not targets (so vary *y_A_* with *x* = 1).

If we are distributing between targets, a single antigenic change is required to escape vaccination at a given target; after this initial change, any subsequent antigenic change at that target will have no effect. Therefore it suffices to consider two possible epidemic sequences, (1) *A*_0_*B*_0_ → *A*_1_*B*_0_ → *A*_1_*B*_1_, and (2) *A*_0_*B*_0_ → *A*_0_*B*_1_ → *A*_1_*B*_1_. Hence

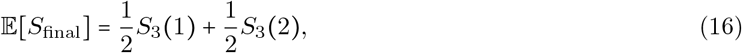

where *S*_3_(*s*) is the remaining individuals without infection acquired immunity following the third epidemic wave for epidemic sequence *s* = 1 or *s* = 2.

If we are instead distributing between variants, since all vaccine doses are directed at target *A* any antigenic change at target *B* will have no effect; moreover, it is possible (depending upon *y_A_*) that two antigenic changes at target *A* will be required to sufficiently deplete the remaining uninfecteds. In combination, this means that *S*_3_(*A*_2_*B*_0_) = *S*_4_(*A*_2_*B*_1_) = ⋯ = *S*_3+*j*_(*A*_2_*B_j_*). Thus there is only one antigenic sequence that matters: *A*_0_*B*_0_ → *A*_1B0_ → *A*_2_*B*_0_, and so 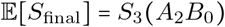.

An important point to note is that when vaccine escape is common, the best outcome we can hope to achieve is 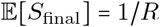. Thus irrespective of vaccine coverage, the population will ultimately be protected by herd immunity acquired through infection. The purpose of vaccination is then to ‘ease’ the population gradually down to the herd immunity threshold so that when it is reached, there are few active infections in the population. In contrast, rapid epidemic dynamics can lead to ‘overshoot’ [1], where many active infections exist when the herd immunity threshold is reached, meaning that although, on average, infections are not able to replace themselves through transmission, overall the susceptible fraction is reduced to below 1/*R*. Consequently, the vaccine strategy that performs best (i.e., maximises 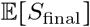) is that which avoids this overshoot. In contrast, when vaccine escape is rare, considerably better outcomes than 1/*R* can be achieved; for example, it is possible under certain conditions for 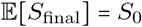.

### 5 Epidemic cases

There are two thresholds which delineate the qualitatively different epidemic regimes discussed in the main text. The first is when the epidemic wave by the initial strain, *A*_0_*B*_0_, reduces the remaining uninfecteds to exactly 1/*R*, that is, *S*_1_ = 1/*R*. To obtain the threshold value of *χ*_0_ for which this occurs (which we denote *T_I/II_*(*p*)), we simultaneously solve equation (12) when *n* = 0 and equation (14) when *n* = 1 and *S*_1_ = 1/*R*, that is, we solve

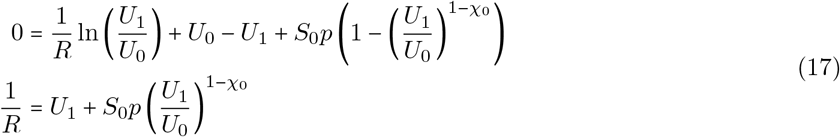

for *U*_1_ and *χ*_0_, then set *χ*_0_ = *T_I/II_*(*p*), which yields

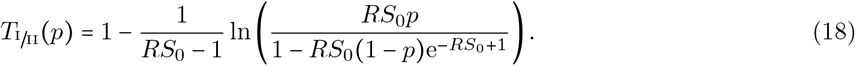

The second threshold occurs when the epidemic wave by the initial strain *A*_0_*B*_0_ is blocked, 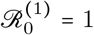. To obtain the threshold value of *χ*_0_ for which this occurs (which we denote *T_II/III_*(*p*)), we solve

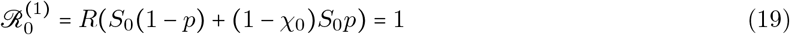

for *χ*_0_ and set *χ*_0_ = *T_II/III_*(*p*) which yields

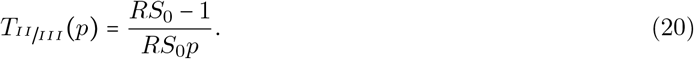

### 6 Metrics to evaluate effectiveness of vaccination

Because vaccination decreases the likelihood of infection, we are primarily interested in reducing the total number of infections over the course of the epidemic. Thus the ‘optimal’ distribution of vaccines across targets and variants is that which maximises 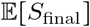. Therefore in the main text we compare the remaining uninfecteds from either distributing across variants, 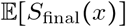, or distributing across targets, 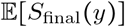, to conventional vaccination, 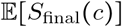, that is, we focus upon 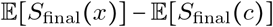 and 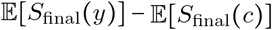; when these quantities are positive, then distributing across targets or variants outperforms conventional vaccination.

Another metric we may be interested in is vaccine efficacy, denoted VE. Specifically, we define

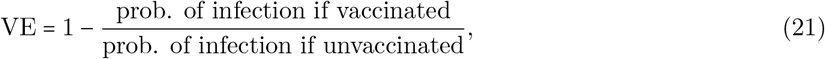

where the probability of infection is calculated over the entire epidemic. Clearly, a higher vaccine efficacy is desirable. If vaccine escape is common, the vaccine efficacy of conventional vaccination is

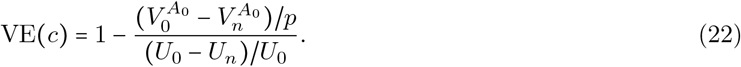

If instead vaccine escape is rare, then there are two possible epidemic sequences of interest: (1) *A*_0_*B*_0_ → *A*_1_*B*_0_ and (2) *A*_0_*B*_0_ → *A*_0_*B*_1_. Let 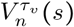 denote the density of individuals vaccinated against *τ_υ_* following the *n*^th^ epidemic of sequence *s* = 1,2. Then the expected vaccine efficacy of conventional vaccination is

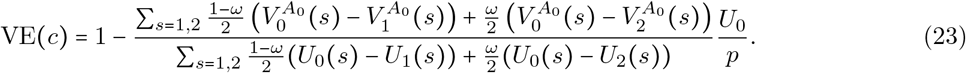

A third metric is what we refer to as vaccine matching, denoted VM. Specifically, given a vaccinated individual is infected, what is the probability they were infected by the target/variant that they were vaccinated against (the ‘matched’ strain)? We calculate this as

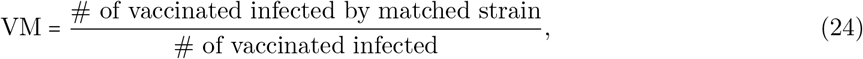

In general, having a higher VM is desirable because although we have assumed the primary effect of vaccination is to reduce the probability of infection, vaccines often have other effects, such as reducing severity of disease. Our model makes no assumptions about what happens to individuals following infection clearance. Thus although it is possible that individuals fully recover from infection, it is also possible that they suffer long-term health consequences or die due to disease complications. Individuals infected by the strain they were vaccinated against are less likely to suffer from long-term health complications or death. Therefore a higher VM indicates a lower individual risk and is beneficial.

For conventional vaccination, if vaccine escape is common, then the only epidemic sequence that matters is *A*_0_*B*_0_ → ⋯ → *A*_1_*B_k_* where *A*_1_*B_k_* is the first strain with variant 1 at target *A*. Thus we can treat this epidemic sequence as *A*_0_*B*_0_ → *A*_1_*B*_0_ → ⋯, and so

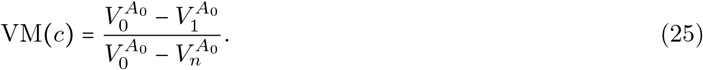

If vaccine escape is rare, then as before the two epidemic sequences of interest are (1) *A*_0_*B*_0_ → *A*_1_*B*_0_ and (2) *A*_0_*B*_0_ → *A*_0_*B*_1_. Therefore

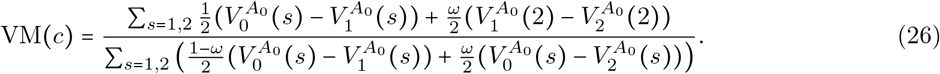

### 7 Vaccines distributed across targets

When we distribute vaccines across targets, there are two sets of sequences of antigenic changes of interest, *A*_0_*B*_0_ → *A*_1_*B*_0_ → ⋯ and *A*_0_*B*_0_ → *A*_0_*B*_1_ → ⋯ (recall that cross-protection is assumed to be limited). The optimal vaccination strategy, *x*\*, is to distribute vaccine doses so as to maximise protection against one of these sequences (Fig. 1). Without loss of generality, focus upon the set of sequences in which the first antigenic change occurs at target A, that is, *A*_0_*B*_0_ → *A*_1_*B*_0_ → ⋯.

When vaccine escape is rare, this means choosing *x* = *x*\* to block the second epidemic wave by *A*_1_*B*_0_, that is, *x*\* satisfies 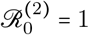. Therefore we solve equation (12) with *n* = 0 for *U*_1_, that is, we solve

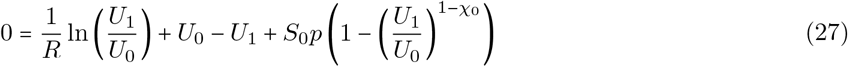

for *U*_1_ and then use that value in

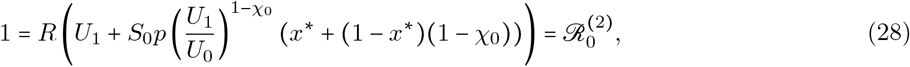

to solve for *x*\*. Solving the second expression for *x*\* as a function of *U*_1_ gives

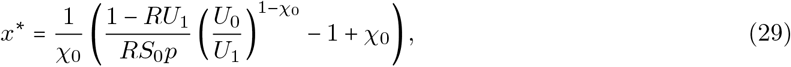

while the first equation can be numerically solved for *U*_1_ (there is no analytic solution of *U*_1_ in this case). Since our choice of *x* does not affect *U*_1_ or *S*_1_, it follows from equation (15) that *x*\* will be independent of *ω*. Of course if *x*\* is a solution then so too is 1 − *x*\*; Fig. 1).

If instead vaccine escape is common (and so epidemics are limited by infection-acquired immunity), then we want to choose *x*\* to satisfy *S*_2_(*A*_1_*B*_0_) = 1/*R*. Therefore we solve

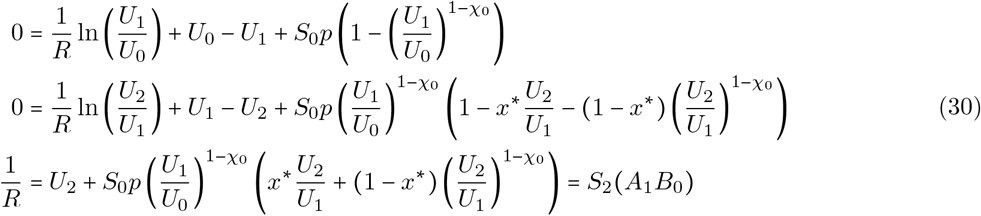

for *x*\*, *U*_1_, and *U*_2_. Note that if we are in case III, that is, *χ*_0_ > *T_II/III_*(*p*), then *U*_1_ = *U*_0_ = *S*_0_(1 − *p*), and we only need to solve for *x*\* and *U*_2_. As before, if *x*\* is optimal, then so is 1 − *x*\* (Fig. 1).

It is straightforward to calculate vaccine efficacy, VE(*x*), when distributing across targets. When vaccine escape is common, there are two sequences of interest: (1) *A*_0_*B*_0_ → *A*_1_*B*_0_ → ⋯ → *A_k_B*_1_ and (2) *A*_0_*B*_0_ → *A*_0_*B*_1_ → ⋯ → *A*_1_*B_k_*, where *A*_1_*B_k_* is the first strain with variant *A*_1_ and *A_k_B*_1_ is the first strain with variant *B*_1_. Thus we can write these sequences as (1) *A*_0_*B*_0_ → *A*_1_*B*_0_ → *A*_1_*B*_1_ and (2) *A*_0_*B*_0_ → *A*_0_*B*_1_ → *A*_1_*B*_1_. Then

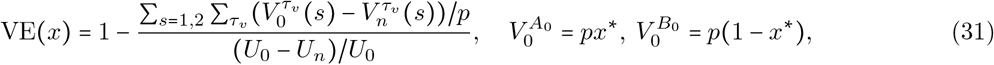

whereas if vaccine escape is rare, there are two possible epidemic sequences, (1) *A*_0_*B*_0_ → *A*_1_*B*_0_ and (2) *A*_0_ *B*_0_ → *A*_0_ *B*_1_, and so

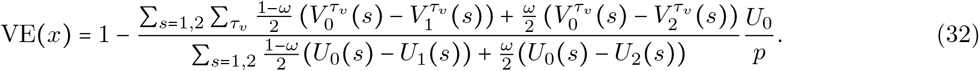

Similarly, we can calculate vaccine matching, VM(*x*), when distributing across targets. When vaccine escape is common, then

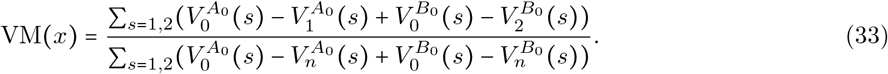

whereas when vaccine escape is rare,

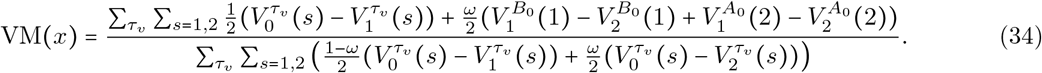

### 8 Vaccines distributed across variants

When we distribute vaccines across variants, the two scenarios of vaccine escape are slightly different. First, for epidemics in which vaccine escape is rare, there are two candidate solutions, *y_A_* = *y*_*A*_0__, in which we maximise protection against the primary variant at target *A*, and *y_A_* = *y*_*A*_1__, in which we maximise protection against the secondary variant at target *A*. Specifically,

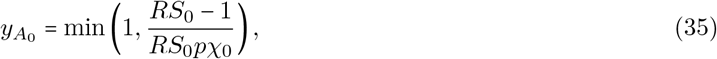

that is, in case II, *y*_*A*_0__ = 1, while in case III, since *U*_1_ = *U*_0_, *y*_*A*_0__ is the solution of

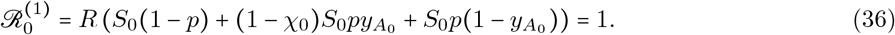

In this strategy, any excess doses not needed to block the *A*_0_*B*_0_ epidemic wave are diverted to protect against the (rare) possibility that a strain carrying the variant *A*_1_ emerges (Fig. 2).

When we maximise protection against the secondary variant, *y_A_* = *y*_*A*_1__, this means choosing *y*_*A*_1__ to (2) satisfy 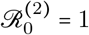, given the next strain is *A*_1_*B*_0_. That is, we solve

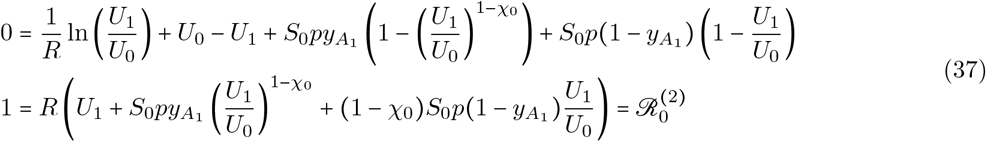

for *y*_*A*_1__ and *U*_1_. Which of *y*_*A*_1__ or *y*_*A*_0__ performs the best will depend upon *R* and the likelihood of vaccine escape, *ω*; when *ω* is small, then *y*_*A*_0__ is best.

Second, for epidemics in which vaccine escape is common, we can always choose *y_A_* to ensure 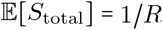. Thus in contrast to distributing vaccine doses between targets, when distributing across variants we can, in theory, always attain the optimum, 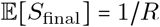. First, we can choose *y_A_* = *y*\* such that after the first epidemic *S*_1_ = 1/*R*. To do so, we solve

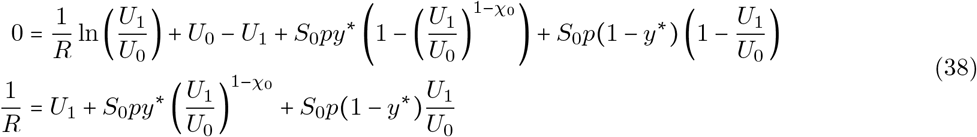

for *U*_1_ and *y*\*, doing so gives

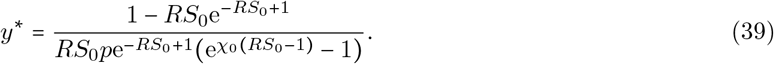

Second, we can choose *y_A_* = *y*^•^ such that *S*_2_ = 1/*R*, if we assume the first antigenic change is *A*_1_*B*_0_ (note that this is justified since *U*_1_ = *U*_2_ =… = *U_j_* for *A*_0_*B_j_*, and so we can ignore all antigenic changes to target *B*). Therefore, we solve

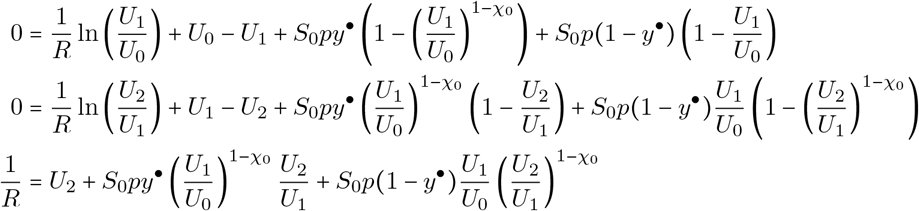

for *U*_1_, *U*_2_ and *y*^•^. Although both *y*\* and *y*^•^ maximise 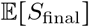, from a public health perspective *y*^•^ is the superior option as it maximises 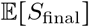 after at least one antigenic change, and so two waves, rather than one epidemic and no antigenic change.

It is straightforward to calculate vaccine efficacy, VE(*y*), when distributing across variants. When vaccine escape is common, there is one sequence of interest, *A*_0_*B*_0_ → ⋯ → *A*_1_*B_k_* → ⋯ → *A*_2_*B_k_*, which we can write as *A*_0_*B*_0_ → *A*_1_*B*_0_ → *A*_2_*B*_0_. Thus

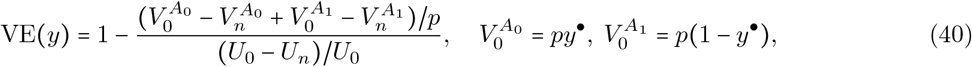

whereas if vaccine escape is rare, there are two possible epidemic sequences, (1) *A*_0_*B*_0_ → *A*_1_*B*_0_ and (2) *A*_0_ *B*_0_ → *A*_0_*B*_1_, and so

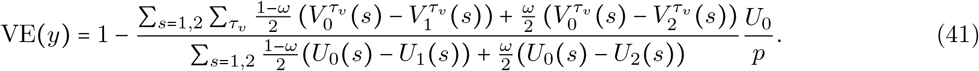

Similarly, we can calculate vaccine matching, VM(*y*), when distributing across variants. When vaccine escape is common, then

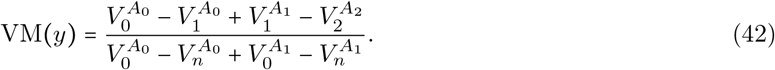

whereas when vaccine escape is rare

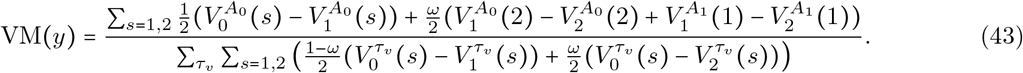

### 9 Vaccines distributed across targets and variants simultaneously

Finally, consider distributing vaccines across both targets and variants. When we are free to choose all three of (*x, y_A_, y_B_*), many different combinations can lead to similar *S_n_*. For example, when antigenic change is not limiting, we have already shown that when only manipulating *y_A_* (with *x* = 1) there are two values of *y_A_* which ensure the optimal outcome, 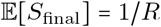. Therefore we have the opportunity to apply further constraints to identify optimal combinations. One approach would be to apply additional metrics. For example, in addition to maximising *S_n_* we could also seek to maximise the average *n* at which *S_n_* is first equal to 1/*R* (this was what separated *y*^•^ from *y*\*), or minimise the variance between epidemic sequences (when *x* = 1/2 and *y_A_* = *y_B_*). Alternatively, we may impose the constraint that we are free to distribute across targets (*x*) and then across both variants in the same way; that is, we take *y_A_* = *y_B_* = *y* and so we are choosing the pair (*x, y*) rather than the triplet (*x, y_A_, y_B_*). The motivation for imposing this constraint is that doing so allows us to see when either distribution dimension has primacy. Specifically, when distributing between targets is more important, then the optimal (*x, y*) will show variation between targets as coverage (*p* and *χ*_0_) changes, while all the doses will be allocated towards the primary variant, *y* = 1. In particular, we expect to see optimal values *x*\* and 1 − *x*\* where *x*\* ≠ 1/2, as observed when distributing across targets only. When distributing between variants is more important, then we should expect *y* to vary with vaccine coverage while doses will be evenly allocated across targets, *x* = 1/2.

The results are much as we would expect (Fig. 3). When the probability of antigenic change is low (*ω* small), it is best to distribute between targets while focusing upon the primary antigen, that is, choose *y* = 1 and *x* = *x*\* where *x*\* is provided in equation (29). As the likelihood of antigenic change increases, however, it becomes optimal to distribute doses between variants, that is, *y* < 1, with doses equally allocated between targets, *x* = 1/2. Specifically, if allocating between variants is the preferred strategy (and so *x* = 1/2), then the optimal choice of *y* will block the next epidemic wave by either strain *A*_1_*B*_0_ or *A*_0_*B*_1_, that is, *y* satisfies 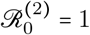 (since *x* = 1/2, 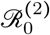 is not sensitive to which strain emerges). Therefore we solve

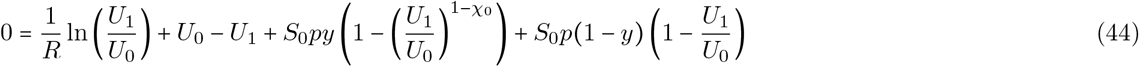

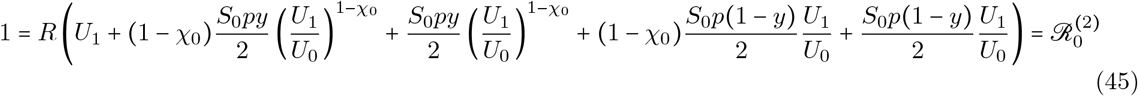

for *U*_1_ and *y* (note the 1/2 multiplier is because it is equally likely that strain *A*_1_*B*_0_ or *A*_0_*B*_1_ will emerge). This can be done with a computer algebra package, but as the answer can only be expressed in terms of the Lambert W function, we do not show it here. Note that as we approach the threshold between case *i* and II, that is, *T_I/II_*(*p*), the solution of (44) in terms of *y* will exceed 1; in this case the optimal solution is *y* = 1 and we resort to varying *x* (i.e., choosing *x*\* from equation (29); Fig. 3).

### 10 Mutations, cross-protection, and multi-strain epidemics

The preceding analysis made three key assumptions. First, antigenic change was assumed to arise without consideration of its source. This is reasonable if antigenic novelty originates outside the focal population. For example, it may come from a reservoir animal population or be otherwise imported from a source population (e.g., influenza A H3N2 from southeast Asia [2, 3]). If instead we focused strictly on antigenic change from *de novo* mutation in the focal population, then unless mutations are very likely, conventional vaccination tends to perform better than our results show; in this case it is typically better to ‘hit hard’ in the hopes of preventing antigenic change, rather than to distribute vaccine doses in anticipation of an unlikely mutation. This is particularly true in case III. The other complicating factor when explicitly considering mutation is whether or not vaccination can induce within-host evolution, biasing which target is most likely to generate novel variants. Although consideration of this possibility is beyond the scope of the current work, in at least some relevant diseases (e.g., influenza A), modelling suggests within-host evolution due to vaccination plays a limited role [4–6].

Second, we assumed cross-protection was limited. The shape, or ‘broadness’ (i.e., how slow the decay of *χ_z_* is for increasing *z*), of the cross-protection function is of no consequence in case I. In cases II and III, however, the primary role of the broadness of vaccine cross-immunity is that it determines the reduction in vaccine protection against new strains. If vaccine cross-immunity is broad, antigenic change will cause a small reduction in vaccine protection, and so a smaller subsequent wave. If, however, cross-immunity is narrow, antigenic change will cause a large reduction in vaccine protection, and so potentially a large wave. Thus increasing cross-protection increases the efficacy of conventional vaccination in case II and III. For example, if antigenic change is not limiting, conventional vaccination is more likely to achieve the optimum of 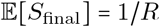 with broad cross-immunity, whereas with narrow cross-immunity, substantial overshoot is more likely. In our model, the shape of cross-immunity will ultimately depend upon the units of antigenic space. For example, if *k_i_* and *k*_*i*+1_ differ by a single, nearly-neutral mutation, cross-protection will tend to be broad. If instead antigenic variation arises in a source population with pre-existing immunity, large differences between *k_i_* and *k*_*i*+1_ may be necessary to escape extinction in the source population, and so cross-protection may be narrow.

Third, we assumed each wave consists of a single strain. Although this may be reasonable if antigenic variation originates elsewhere, it is less likely when variation is generated by *de novo* mutation. If our results were extended to include multi-strain waves, the predicted efficacy of conventional vaccination would be weakened, since whenever an escape mutant arises during an ongoing wave, there are more individuals without infection-acquired immunity and so available to be infected by the escape mutant. This would lead to a larger wave by the strain with limited vaccine protection.

---

**Figure 1:**
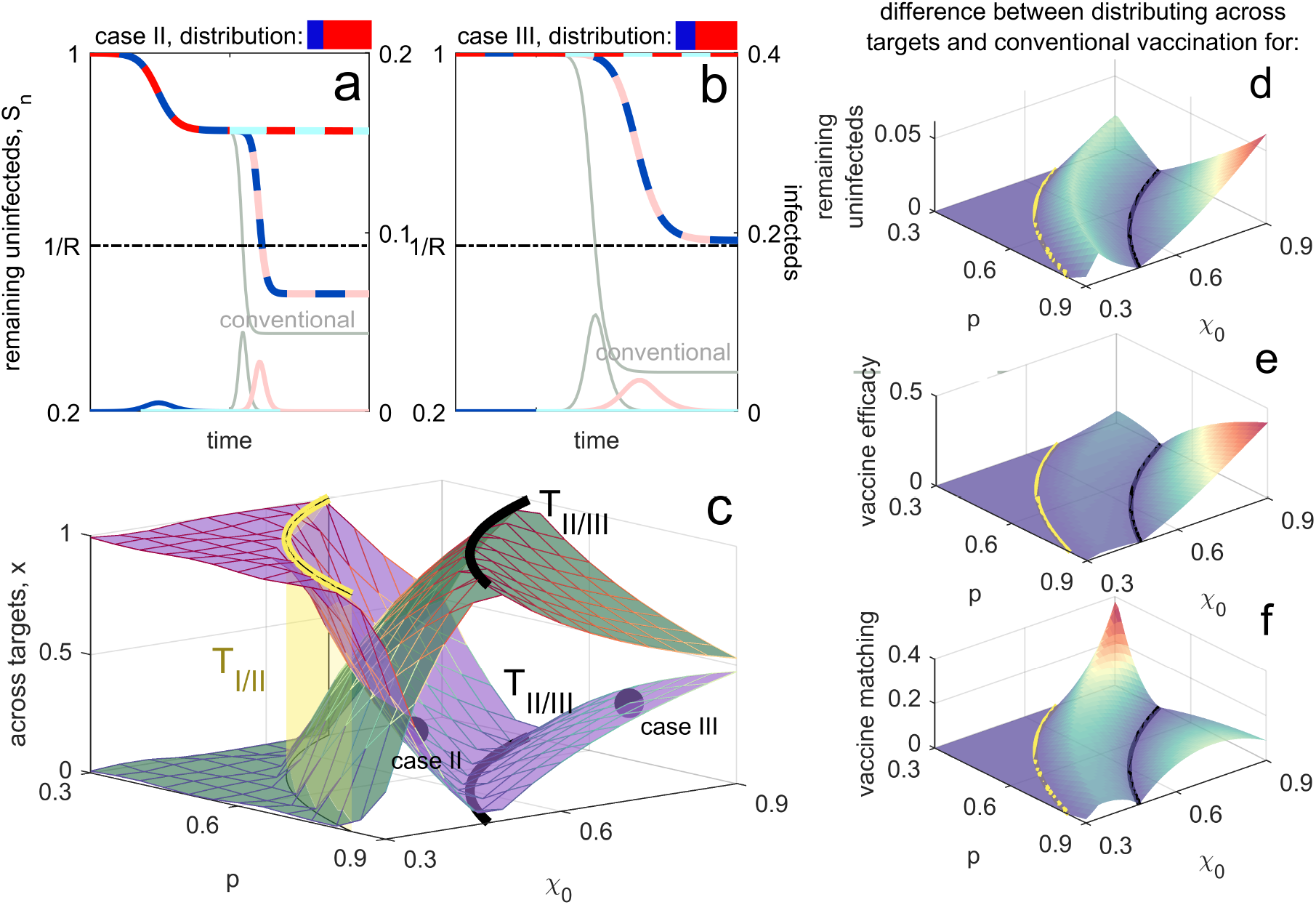
Optimal distribution of vaccines across targets when vaccine escape is rare. Panels **a,b** show example epidemic dynamics for case II (panel **a**) and case III (panel **b**), with conventional vaccination (light grey) shown for reference. Note that following the initial epidemic wave by strain *A*_0_*B*_0_, there are two possible sequences of interest, *A*_0_*B*_0_ → *A*_1_*B*_0_ and *A*_0_*B*_0_ → *A*_0_*B*_1_; both are shown. The optimal distribution across targets (shown in panel **c** as vaccine coverage, *p*, and strength, *χ*_0_, vary) maximises the remaining susceptibles averaged over these sequences; when *x* is optimal, so is 1 − *x* (purple, green). Panels **d-f** show how distributing across targets outperforms conventional vaccination for three metrics; positive values indicate the degree to which mosaic vaccination is superior. Panel **d**, shows the difference in remaining uninfecteds, 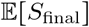. Panel **e** shows the difference in vaccine efficacy, measured as attack rate over the course of the epidemic. Panel **f** shows the difference in vaccine matching, defined as the probability that an individual that is both vaccinated and infected is infected by a strain that they were vaccinated against. In all panels, the thresholds *T_I/II_* (yellow surface) and *T_I/II_* (black lines) are included for reference; beyond the yellow surface, vaccine efficacy is sufficiently low that evolution has no effect on the epidemic, and any distribution of vaccines between targets will produce the same outcome (for a given *p* and *χ*_0_).

**Figure 2:**
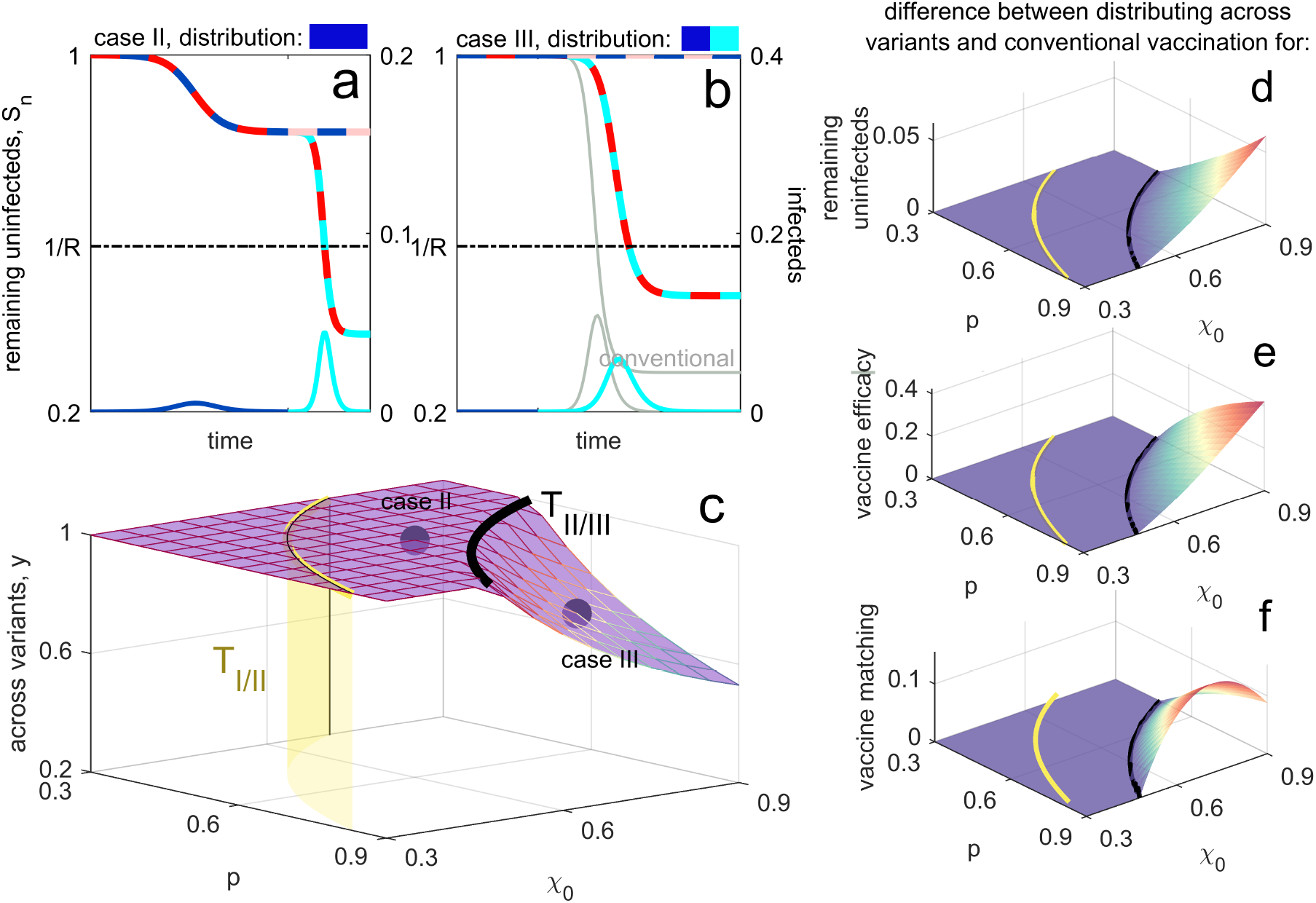
Optimal distribution of vaccines across variants when vaccine escape is rare. Panels **a,b** show example epidemic dynamics for case II (panel **a**) and case III (panel **b**), with conventional vaccination (light grey) shown for reference (in panel **a**, conventional vaccination is same as distributing between variants). Following the initial epidemic wave by strain *A*_0_*B*_0_, there are two possible sequences of interest, *A*_0_*B*_0_ → *A*_1_*B*_0_ and *A*_0_*B*_0_ → *A*_0_*B*_1_; both are shown. The optimal distribution across variants (shown in panel **c** as vaccine coverage, *p*, and strength, *χ*_0_, vary) maximises the remaining susceptibles averaged over these sequences. Panels **d-f** show how distributing across variants outperforms conventional vaccination for three metrics; positive values indicate the degree to which mosaic vaccination is superior. Panel **d**, shows the difference in remaining uninfecteds, 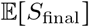. Panel **e** shows the difference in vaccine efficacy, measured as attack rate over the course of the epidemic. Panel **f** shows the difference in vaccine matching, defined as the probability that an individual that is both vaccinated and infected is infected by a strain that they were vaccinated against. In all panels, the thresholds *T_I/II_* (yellow surface) and *T_I/III_* (black lines) are included for reference; beyond the yellow surface, vaccine efficacy is sufficiently low that evolution has no effect on the epidemic, and any distribution of vaccines between targets will produce the same outcome (for a given *p* and *χ*_0_).

**Figure 3:**
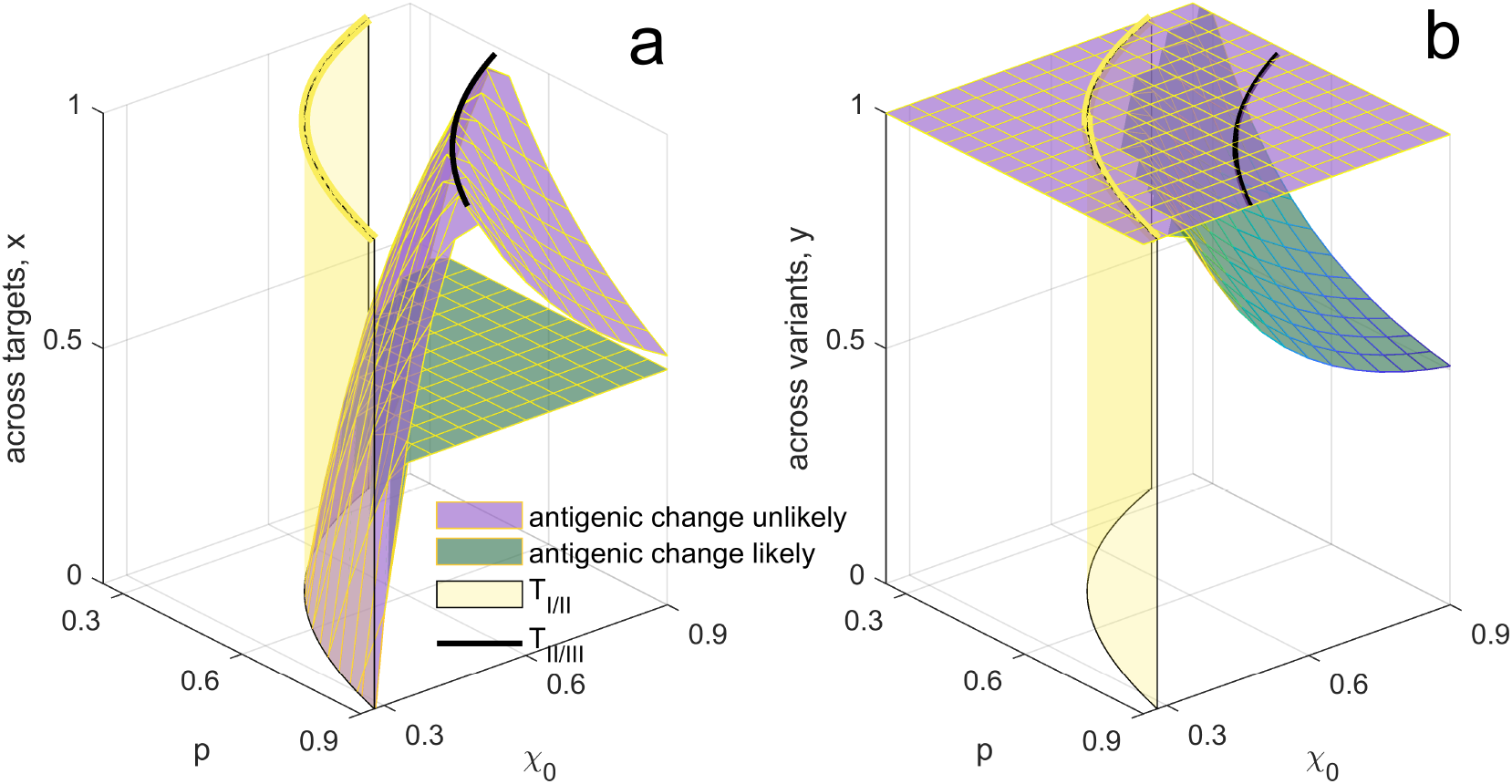
Optimal distribution of vaccines between targets (panel **a**) and variants (panel **b**). In both panels, the purple surface corresponds to the optimal distribution when antigenic change is unlikely (i.e., *ω* is small) while the green surface corresponds to the optimal distribution when antigenic change is likely. As before, beyond the yellow surface, vaccine efficacy is sufficiently low that evolution has no effect on the epidemic outcome. In general, when antigenic change is unlikely (purple surfaces) we should distribute between targets and protect against primary variant (*y* = 1), whereas when antigenic change is likely (green surfaces) we should distribute between variants with doses evenly allocated between targets (*x* = 1/2). Note that for visual clarity in panel **a**, we have only shown one possible optimal solution; but if *x* is optimal, then so is 1 − *x*.

## Notes

### Competing Interest Statement

The authors have declared no competing interest.

